# CXCR3 on transferred T cells enables vaccine-boosted adoptive T-cell therapy in solid tumors

**DOI:** 10.64898/2026.10.04.753375

**Authors:** Vinnycius Pereira Almeida, Lucy C. Garner, Sara Abdel Malak, Amanda Wicki, James McAuliffe, Laurine Noblecourt, Silvia Panetti, Emily E. Steffke, Aneesha Bhandari, Lyn Yong, Ali Amini, Claire Hutchings, Francesco Andreata, Yuanqin Yang, Audrey Gérard, Matteo Iannacone, Ricardo A. Fernandes, Paul Klenerman, Benoit Van den Eynde, Carol Sze Ki Leung

**Author notes:** These authors contributed equally to this work (Lucy C. Garner, Sara Abdel Malak).

## Abstract

Adoptive T-cell therapy (ACT) has demonstrated marked clinical benefit in hematological malignancies but has limited efficacy against solid tumors. Cancer vaccines can improve ACT, but the mechanisms underlying this combination are poorly understood. Here, we combined ACT with chimpanzee adenovirus ChAdOx1 (ChAd) and modified vaccinia Ankara (MVA) encoding tumor antigens. Whereas PD1 blockade failed to improve ACT, the combination with ChAd/MVA vaccination drove durable regression of established tumors, correlating with expansion of tumor-reactive T cells in lymphoid organs and increased tumor infiltration by T cells. Mechanistically, ChAd priming induced CXCR3 ligands (CXCL10 and CXCL9) in the vaccine-draining lymph node, thereby recruiting transferred T cells to this node independently of cognate antigen. CXCR3 expression on the transferred T cells, but not on host cells, was required for the anti-tumor effect of the combination therapy and favored T-cell accumulation in the vaccine-draining lymph node and the tumor. These findings identify CXCR3 on the transferred T cells as a determinant of ACT plus vaccination synergy and provide a rationale for combining viral-vector vaccines with TCR-T therapies, and for selecting or engineering T cells to retain CXCR3 expression.

**One Sentence Summary:** Viral vector vaccines synergize with TCR-T cell therapy to control solid tumors in a CXCR3-dependent manner.

## INTRODUCTION

Systemic infusion of cytotoxic tumor-specific T cells to control neoplasia (adoptive T-cell therapy, ACT) has demonstrated good clinical efficacy in hematological malignancies but has limited efficacy against solid tumors in humans (*1*). Factors contributing to therapy failure include insufficient T-cell numbers, suboptimal in vivo T-cell expansion, T-cell exhaustion, poor immunological memory formation, low cell persistence, limited tumor infiltration and suboptimal targets (*2, 3*). Nevertheless, technological and methodological advancements culminated in the approval of the first ACTs against solid tumors – lifileucel and afamitresgene autoleucel – by the US Food and Drug Administration in 2024. The approved T-cell products were either expanded from the patients’ tumors, the tumor-infiltrating lymphocytes (TILs) (lifileucel), or from T cells collected from blood and genetically modified to express a TCR against the tumor antigen MAGE-A4 in HLA-A*02 (afamitresgene autoleucel) (*4, 5*). While these therapies have demonstrated clinically meaningful responses and achieved regulatory approval, only a subset of patients experience durable benefit, highlighting the need for new strategies to further improve the efficacy of ACT against solid tumors.

A central determinant of ACT efficacy is the differentiation state of the transferred T cells. This state is shaped by the initial T-cell priming and subsequent interactions with other cells (e.g. dendritic cells, DCs) and soluble factors (e.g. cytokines, such as IL2) (*6*). Exposure to appropriate levels of stimulation enables T cells to acquire potent cytotoxic activity (*7, 8*) against tumor cells while retaining the ability to acquire memory capacity.Conversely, inadequate T-cell activation or overstimulation can lead to T-cell exhaustion, resulting in poor tumor response despite the presence of anti-tumor effector T cells (*9, 10*). Evidence in mice and humans showed that less differentiated T cells, such as central memory (T_CM_; CD62L+ CD44+ in mice and CD45RA-CCR7+ CD62L+ in humans) and stem-like (TCF1+ PD1+) T cells, result in better anti-tumor responses than effector T cells when used for ACT (*3, 11–14*), particularly when combined with vaccines (*3*).

Cancer vaccines can amplify the anti-tumor activity of ACT in vivo (*15–19*), and this synergy relies on functional secondary lymphoid tissues (*3*). ACT has been combined with dendritic-cell (*20*), peptide (*17, 19*), viral-vector (*19, 21*), and mRNA (*22*) vaccines, with mixed anti-tumor outcomes, often relying on lymphodepleting preconditioning and IL2 supplementation, which increase treatment burden and side effects (*1*). For example, targeting the gp100 antigen in an ACT plus vaccine setting was ineffective in metastatic melanoma (*19*), whereas vaccine-primed T-cell therapy combined with a DC vaccine reached a median overall survival of 14.2 months in ovarian cancer, against a historical median of 6.2 months for fourth-line chemotherapy (*15*). Although it is well established that CD62L and CCR7 expression is required for homing of T cells to lymphoid tissues, how different vaccination regimens act within these tissues to expand the transferred tumor-reactive T cells, and whether this requires cognate antigen, remains poorly defined.

To identify novel therapeutic approaches to improve ACT against solid tumors and to define the determinants of ACT plus vaccines efficacy, we used the replication-incompetent viral vectors chimpanzee adenovirus (ChAdOx1 or ChAd) and modified vaccinia Ankara (MVA) poxvirus viral vectors. These vaccines mimic viral infection, inducing strong cytotoxic CD8+ T-cell responses even against poorly immunogenic antigens (*23, 24*). These vectors are clinically validated, safe (*25, 26*), immunogenic and have shown clinical promise against prostate cancer antigens (*26*). In mice, ChAd/MVA targeting the MAGE-type tumor antigen P1A showed therapeutic efficacy when combined with the immune checkpoint inhibitor (ICI) αPD1 (*27*). Optimal ChAd immunogenicity relies on intact STING and IFNAR (*28*) and the activation of mucosal-associated invariant T (MAIT) cells by the induction of cytokines, such as IFNα and IL18 (*29*). Furthermore, ChAd/MVA vaccines promote a more inflammatory tumor microenvironment (TME), with increased levels of *Cxcl9*, *Cxcl10* and *Ccl5*, and a high magnitude of tumor-specific CD8+ T-cell responses (*27, 30*). These properties make ChAd/MVA heterologous prime boost a promising tool to improve ACT anti-tumor effects. We therefore combined ACT with ChAd/MVA vaccines and defined the cellular and chemokine determinants of their synergy, identifying CXCR3-dependent T-cell trafficking as the critical axis.

## RESULTS

### Viral vector vaccination improves anti-tumor efficacy of adoptive T-cell therapy

For ACT, we used CD8+ T cells from the spleen of TCRP1A mice (*31*), which recognize the H-2L^d^-restricted dominant epitope of P1A (P1A_35-43,_ LPYLGWLVF), a shared non-mutated tumor-specific MAGE-type antigen (*32*). We activated T cells in vitro by co-culture with tumor cells engineered to express P1A and B7-1 (fig. S1A). After 4 days, the expanded T cells expressed memory (CD127) and activation (CXCR3, PD1, CD44) markers while preserving mostly a T_CM_ phenotype (≈ 74% CD62L+ CD44+) (fig. S1B).

We injected mice bearing established (∼150mm^3^) P1A-expressing 15V4T3 mastocytoma tumors with 1-2E+5 activated TCRP1A CD8+ T cells i.v., either alone or combined with αPD1 or ChAd-P1A and MVA-P1A vaccines (Fig. 1A). ACT alone showed some efficacy to reduce tumor growth and increase survival (Fig. 1, B and C). While ACT efficacy was not improved by adding 3 injections of αPD1, it was dramatically increased upon combination with the vaccines (without preconditioning and IL2 supplementation) (Fig. 1, B and C), with complete tumor-rejection observed in 10 of 13 mice, without signs of toxicity (weight loss) (Fig. 1C and fig. S1C).

**Fig. 1.**
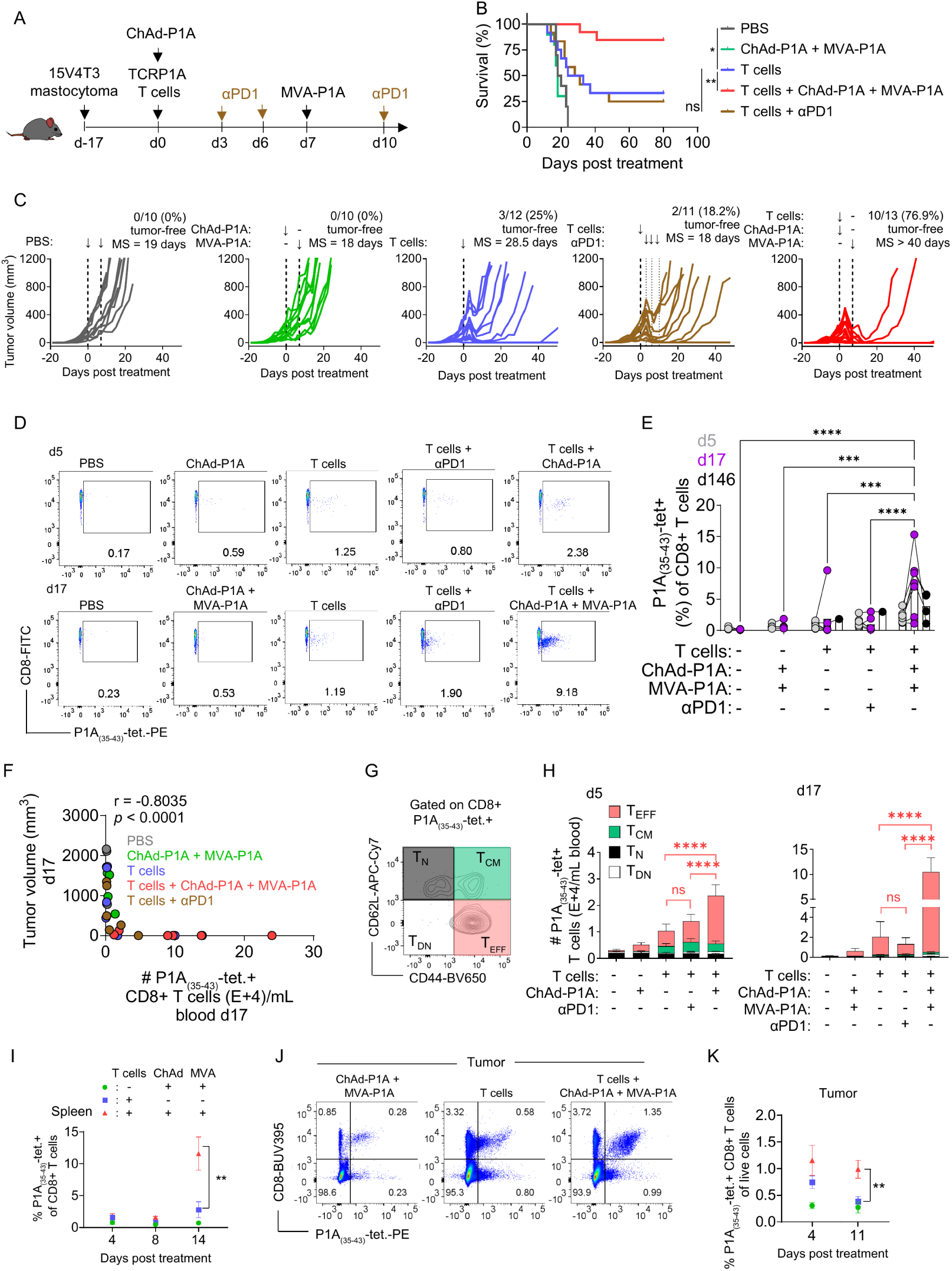
Combination immunotherapy with P1A T cells and viral vector vaccination results in complete and durable tumor control. (A) Treatment scheme. (B) Kaplan-Meier survival curve after combination therapy and monotherapy controls; statistical analysis by log-rank (Mantel-Cox) test. (C) Individual tumor growth curves over time and median survival (MS). Data in B and C were pooled from two independent experiments. (D) Representative plots and (E) percentage of P1A-specific (LPYLGWLVF–H-2Lᵈ tetramer⁺) cells out of CD8⁺ T cells, 5, 17 and 146 days after ACT. Statistical analysis was performed on log-transformed frequencies using a mixed-effects model, followed by Tukey’s multiple-comparisons test; the model was fit on days 5 and 17, and day 146 (survivors only) is shown descriptively. (F) Correlation between P1A-specific CD8⁺ T cells and tumor volume on day 17, assessed by two-tailed nonparametric Spearman correlation. (G) Representative plot of CD62L and CD44 populations gated on P1A-specific T cells and (H) their quantification on days 5 and 17 post-treatment, analyzed by repeated-measures two-way ANOVA with Tukey’s multiple-comparisons test. (I) Percentage of tetramer⁺ cells out of CD8⁺ T cells in the spleen on days 4, 8 and 14, pooled from independent experiments. (J) Representative plot of P1A-specific T cells in the tumors under the different treatments on day 11 and (K) their proportion in the TME on days 4 and 11. For I and K, terminal harvests at each timepoint comprise independent groups of mice; data are shown as mean ± SEM and were analyzed separately at each timepoint by Kruskal-Wallis test followed by Dunn’s post-hoc test with Benjamini-Hochberg correction. *p < 0.05, **p < 0.01, ***p < 0.001, ****p < 0.0001.

We monitored the presence of TCRP1A CD8+ T cells in the blood of mice at different timepoints following treatment using H2-L^d^/ P1A(_35-43_) tetramers (Fig. 1, D and E). Low percentages of P1A-specific T cells (<2% of CD8+ T cells) were observed after vaccination or ACT only and were not increased by combining ACT with αPD1. However, combining ACT with ChAd-P1A and MVA-P1A vaccines significantly increased the percentage of P1A-specific T cells at day 17 (>9%) compared to either therapy alone (Fig. 1E). P1A-specific T-cell numbers were inversely correlated with tumor volume (Fig. 1F). Circulating P1A-specific T cells produced IFN*γ* and TNF upon activation (fig. S1D), aligning with an increased number of effector T cells (CD44+CD62L-) in the combination therapy group (Fig. 1, G and H). A similar expansion of P1A-specific T cells was observed in the spleens and the tumors of mice after ACT combined with vaccination (Fig. 1, I to K). Study of tumor T-cell infiltrates at timepoints later than day 11 was not possible due to complete tumor resolution in most mice receiving the combination treatment (fig. S1E). As in the blood, the abundance of P1A-specific T cells in the spleen correlated inversely with the tumor mass (fig. S1E). Interestingly, the vaccination also promoted long-term persistence of the P1A-specific T cells in the blood, spleens, draining lymph nodes (dLN), and bone marrow of surviving mice (fig. S1, F and G).

Altogether, these results show a synergy between ACT and cognate vaccination with ChAd/MVA viral vectors, inducing strong expansion of tumor-specific T cells, leading to superior tumor resolution compared to monotherapeutic approaches.

### The priming injection of ChAdOx1 and MVA boost are required for tumor control

Both ChAd and MVA viral vectors are DNA viruses with intrinsic capacities to stimulate innate immune responses in addition to promoting specific immunity to the recombinant antigens they encode (*28, 33*). To evaluate which vaccines’ properties were sufficient to enhance ACT efficacy against solid tumors, we compared the combinatorial use of viral vectors encoding the relevant tumor antigen P1A, as used before, with the same viral vectors encoding an irrelevant antigen (i.a.) in tumor-bearing mice (Fig. 2A).

**Fig. 2.**
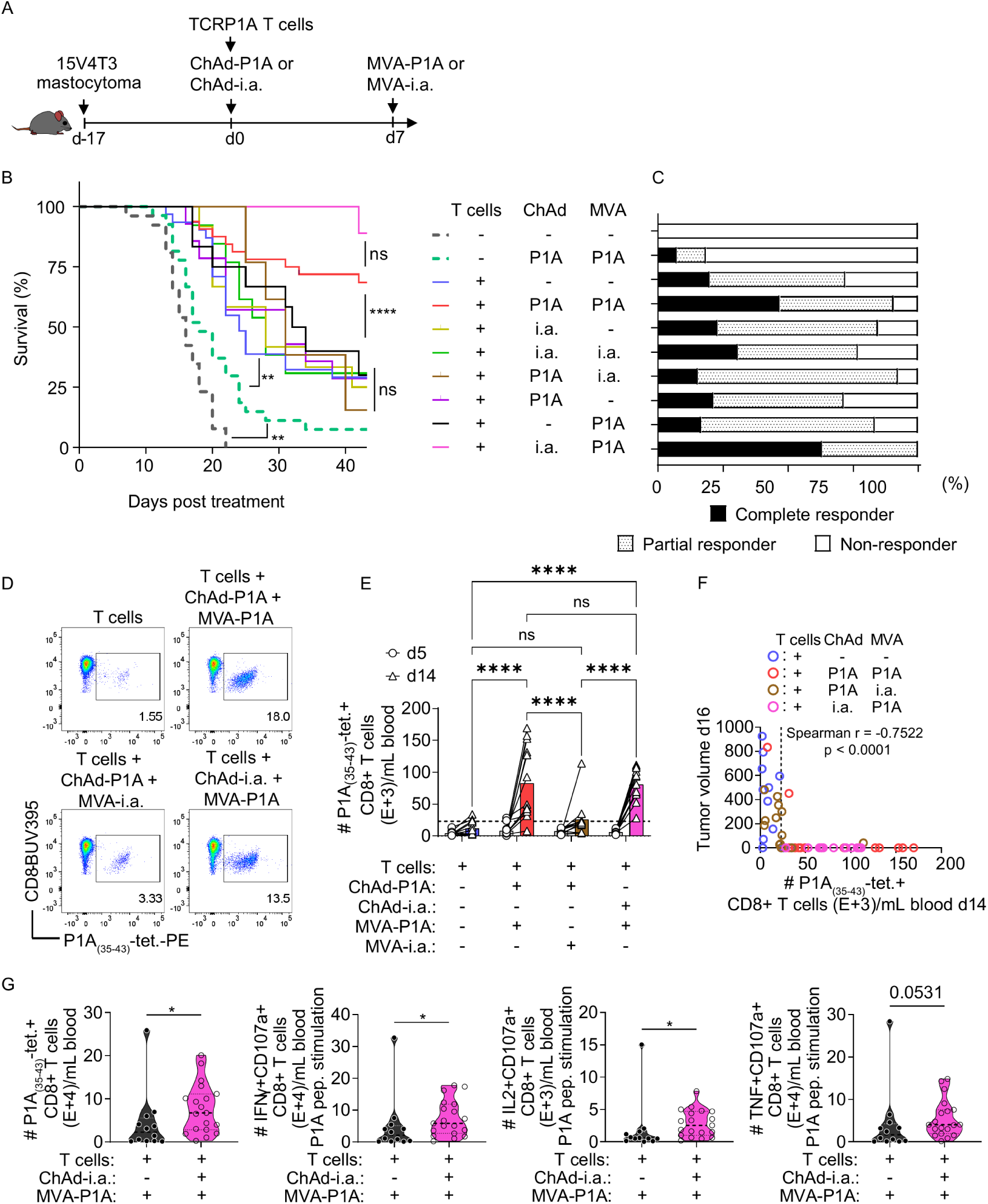
Priming with ChAd expressing an irrelevant antigen is sufficient to significantly improve tumor control by adoptive T-cell therapy when boosted with MVA-P1A. **(A)** Treatment scheme. **(B)** Kaplan-Meier survival curves of the indicated treatment groups. Statistical analysis was performed using log-rank (Mantel-Cox) test. **(C)** Mice in different groups were stratified into non-responders, partial responders and complete responders, which were plotted as a percentage of the total number of mice in each group. **(D)** Representative plots of P1A(_35-43_)-tetramer+ CD8+ T cells under the different treatment groups, which are quantified in (E). **(E)** Bar graphs display circulating P1A-specific CD8+ T cell numbers on days 5 (open circles) and 14 (open triangles) under different experimental conditions; bar color denotes the treatment group. Statistical analysis was performed using the 2way ANOVA method followed by Tukey’s correction for multiple comparisons test. **(F)** Spearman correlation between the number of P1A-specific CD8+ T cells per mL of blood on day 14 and tumor volumes on day 16. Each dot represents a mouse of the corresponding labeled group, and the dotted line marks the 2.3E+4 cells ml⁻¹ threshold. **(G)** Number of tetramer+ CD8+ T cells per mL of blood on days 12-14, as well as number of IFNγ+ CD107a+, IL2+ CD107a+, and TNF+ CD107a+ CD8+ T cells per mL of blood after stimulation with overlapping 15mers P1A peptides.

Priming with ChAd-i.a. followed by a boost with MVA-i.a. did not synergize with ACT, indicating that the presence of the relevant tumor antigen in either vaccine (or both) was required for improving the tumor control and survival (Fig. 2, B and C and fig. S2, A to F). The relevant antigen was specifically required in the MVA, as a ChAd-P1A prime followed by an MVA-i.a. boost also failed to increase tumor control compared to ACT alone (Fig. 2, B and C and fig. S2, G and H). Strikingly, the relevant antigen was not required in the ChAd prime, and a ChAd-i.a. prime followed by an MVA-P1A boost was as efficient as ChAd-P1A/MVA-P1A to improve tumor control upon ACT (Fig. 2, B and C and fig. S2I). Furthermore, the ChAd-i.a. prime followed by MVA-P1A boost induced a strong expansion of P1A-specific CD8+ T cells, which was equal to the expansion observed with the ChAd-P1A/MVA-P1A and correlated with tumor control (Fig. 2, D to F). Such an expansion was not observed when ACT-treated mice were vaccinated with MVA-P1A (same timepoint as usual, day 7) in absence of ChAd prime (Fig. 2G), which correlates with the tumor growth kinetics (fig. S2J), indicating the importance of the ChAd prime to allow the effect of the subsequent MVA-P1A. Interestingly, the order of the vaccines was also crucial as priming with MVA-P1A and boosting with ChAd-P1A in the context of ACT did not increase tumor control compared to ACT monotherapy (fig. S2K).

We observed a similar synergistic effect of ACT in combination with cognate ChAd/MVA in B16F10-OVA, a melanoma model expressing chicken ovalbumin (OVA) as a surrogate tumor antigen (fig. S3, A to G). Consistent with the mastocytoma model, combination of OVA-ACT with ChAd/MVA-OVA resulted in a longer median survival and more complete responders than any of the monotherapies tested against melanoma (fig. S3, B to G). Also, in line with previous results, the MVA encoding the relevant antigen was crucial for the combination therapy efficacy (fig. S3, H and I). However, in contrast to the mastocytoma model, ACT plus ChAd-i.a. prime and MVA-OVA boost was not sufficient for improving mice survival compared to ACT alone (*p=*0.4494, fig. S3, C, F, I, and J), indicating that the effect of ChAd-i.a. depends on the tumor context and/or the target antigen.

To investigate how the different vaccine combinations impacted the endogenous and transferred antigen-specific CD8+ T cells in the context of ACT, we made use of the CD45.1 isoform expressed on the transferred OVA T cells and the CD45.2 expressed on the endogenous host cells (fig. S3K). We observed that the vaccines induced expansion of the transferred (but not the endogenous T cells) on days 5 and 13, as long as the relevant MVA boost was added to the vaccination scheme (fig. S3, L and M), the transferred but not the endogenous T cells being inversely correlated with tumor size (fig. S3, N and O). Previous reports showed that viral vectors indeed promote transferred T-cell expansion, but as soon as day 12 post treatment, the circulating endogenous T cells surpass the transferred ones, becoming predominant in the blood (*34*). However, it is not clear if this pattern is consistent across lymphoid tissues in later timepoints. Thus, we investigated the persistence of the transferred versus endogenous OVA-specific T cells in different tissues after ≈7.5 months in the combination therapy survivors (fig. S3P). Consistent with the early timepoints post vaccination, the combination therapy resulted in higher proportions of transferred OVA-specific T cells in the blood as compared to endogenous T cells. Surprisingly, an opposite trend was observed in the dLN, and a comparable frequency of transferred and endogenous T cells was observed in the spleen and the bone marrow (fig. S3P).

Thus, the synergistic effect of ChAd/MVA with ACT requires a cognate MVA boost, but not necessarily a relevant antigen-coding prime, depending on the tumor type. The effect of the boost is closely associated with the expansion of transferred antigen-specific T cells, which can be detected in lymphoid tissues several months after infusion, suggesting memory formation. Furthermore, ChAd priming is required for the combination therapy to have its optimal anti-tumor effect.

### ChAdOx1 promotes CXCL10 expression in the vaccine-draining lymph nodes

To investigate how ChAd priming influences subsequent treatments, we investigated the early immunological response following ChAd vaccination. In a previous report, we showed that ChAd activates the cGAS-STING pathway and induces high levels of IFNα, which is critical for ChAd immunogenicity (*28*). However, it is not clear whether this pathway is also critical in the context of ACT combined with vaccination in the tumor setting. We profiled cytokines in the serum 6 hours after ChAd i.m. vaccination (Fig. 3, A and B) and observed a striking increase in the levels of CXCL10 and IFNα, with an average fold-change relative to non-vaccinated controls (PBS) of 6.13 and 3.95, respectively (Fig. 3B). We reanalyzed bulk RNA-seq data from human whole blood 24 hours after ChAd vaccination (*35*) and observed a similar upregulation of genes belonging to the cGAS-STING pathway, including the CXCR3 ligands, CXCL9, CXCL10 and CXCL11 (fig. S4A).

**Fig. 3.**
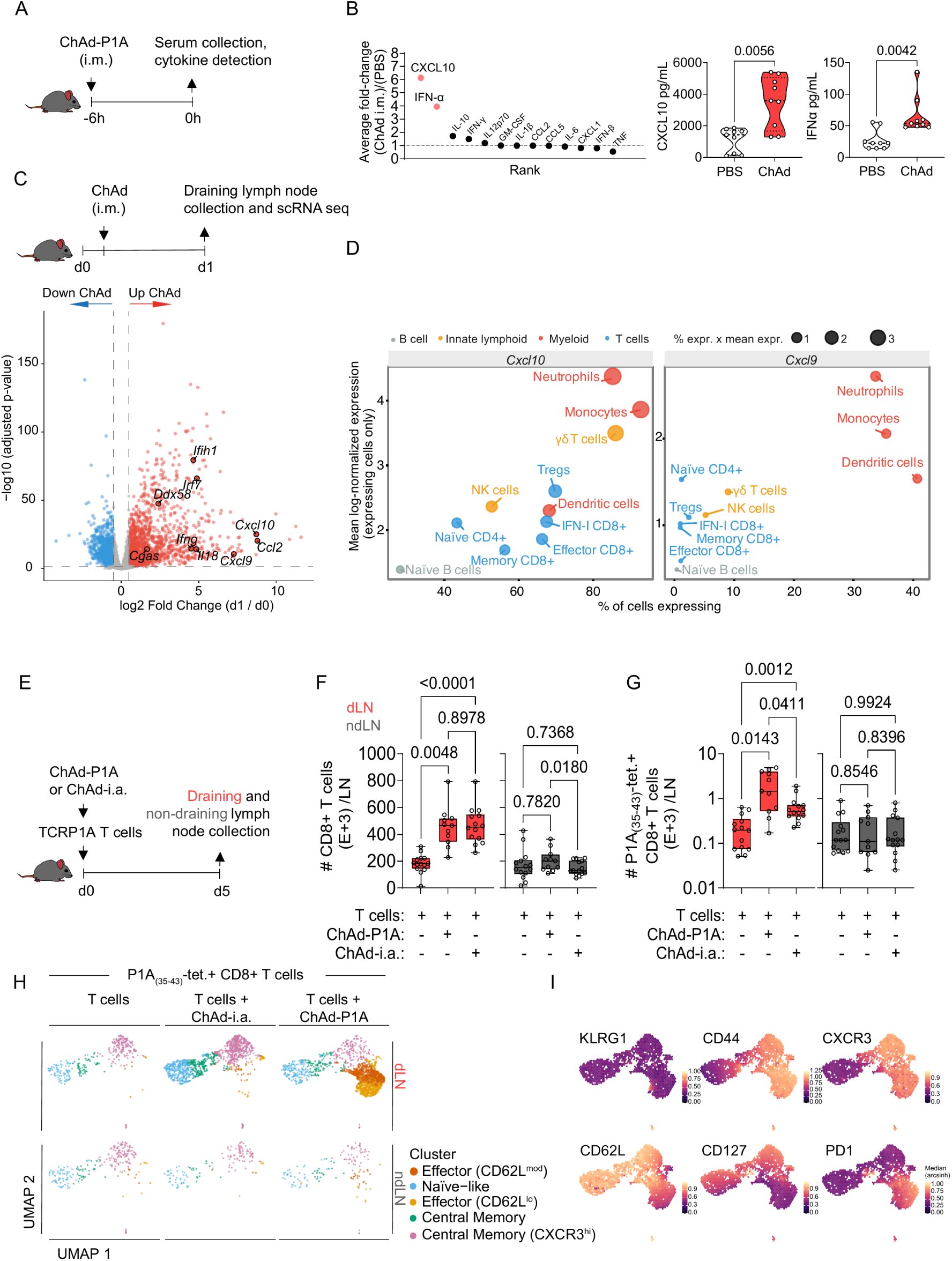
ChAdOx1 induces CXCL10 in the vaccine-draining lymph node and promotes antigen-independent accumulation of antigen-specific CD8⁺ T cells. (A) Mice were vaccinated with ChAd-P1A intramuscularly (i.m.) and 6 hours later serum was collected. (B) Average fold-change of two independent experiments comparing cytokines after ChAd i.m. relative to non-vaccinated (PBS) control. Concentrations of cytokines are shown on the right panel. (C) Mice were vaccinated with ChAdOx1 and 1 day after the vaccine-draining lymph nodes were collected. Single cell RNA sequencing was performed on non-vaccinated (day 0) and vaccinated mice (day 1). Volcano plot showing single-cell RNA-seq data from draining lymph node after ChAd vaccination. Cells (Day 0 and Day 1 post-vaccination) were processed using Seurat. Pseudobulk differential expression (Day 1 vs Day 0) was performed using DESeq2 (n = 5 mice per timepoint; design ∼ day, unpaired). Genes with |log2FC| > 0.5 and padj < 0.05 (Benjamini-Hochberg) were considered differentially expressed. **(D)** Dot plot showing *Cxcl10* (left) and *Cxcl9* (right) expression across single-cell RNA-seq-defined immune cell populations at day 1 post-vaccination. Dot size indicates the percentage of cells expressing the gene x the mean expression within each population; dot color indicates the cell lineage. **(E)** Mice received TCRP1A T cells and ChAd-P1A or ChAd-i.a.; 5 days after, vaccine-draining and non-draining lymph nodes were collected. **(F)** Box plots with total number of CD8+ T cells and (G) total number of P1A(_35-43_)-H-2L^d^-tetramer+ T cells in the lymph nodes. **(H)** Uniform Manifold Approximation and Projection (UMAP) of P1A(_35-43_)-H-2L^d^-tetramer+ T cells phenotypes. Flow cytometry data was exported from FlowJo and processed in R. **(I)** Density plots displaying the per-cell arcsinh-transformed marker expression values of each marker in P1A(_35-43_)-H-2L^d^-tetramer+ CD8+ T cells from the T cells plus ChAd-P1A group.

As ACT combined with either ChAd or MVA alone was not sufficient to reduce tumor burden compared to ACT monotherapy, we investigated how the prime vaccination could affect the cytokine profile of the subsequent boost vaccination (fig. S4B). IFNγ and CXCL10 were highly enhanced in the serum 6 hours after the last dose in ChAd/MVA vaccinated mice compared to MVA alone, indicating that ChAd priming substantially impacts the magnitude of the cytokine response elicited by the MVA boost (fig. S4, C to E). Of note, CXCL10, which was induced 6 hours after a single ChAd prime, returned to normal levels 7 days later in the absence of a boost (fig. S4E).

Cytokines and chemokines are important soluble factors that determine cell profiles and chemotaxis in tissue niches over time (*36*). To investigate whether CXCL10 overexpression was also evident in the vdLN and to characterize the early transcriptional response to ChAd vaccination in this tissue, we performed single-cell RNA-sequencing (scRNA-seq) of the vdLN on day 1 after vaccination (Fig. 3C). Consistent with our serum cytokine profile, *Cxcl10* was among the most strongly expressed genes in the vdLN compared with naïve controls (Fig. 3C). ChAd vaccination also induced genes associated with the cGAS-STING pathway (*Irf7, Cgas*), RIG-I-like receptors (*Ddx58*, *Ifih1*), cytokines (*Ifng*, *Il18*) and chemokines (*Cxcl9*, *Ccl2*) (Fig. 3C). Gene Set Enrichment Analysis (GSEA) demonstrated enrichment of IFNα-, IFNγ- and TNF-signaling pathways following ChAd vaccination, together with antigen processing and cross presentation and oxidative phosphorylation pathways (fig. S4F).

Although the CXCR3 ligands present some degree of redundancy in attracting T cells in infection models in vivo, their regulation is controlled by different promoter elements and cytokine combinations (*36*). CXCL10 is induced in response to inflammatory signals, such as IFNγ, IFNα/β, and TNF (*37–39*), whereas CXCL9 is exclusively induced by IFNγ (which is enhanced in the presence of TNF) (*36*).

Next, we defined the cellular sources of CXCL10 and CXCL9 in the vdLN following ChAd vaccination. Monocytes, neutrophils and γδ T cells exhibited both the highest proportion of *Cxcl10*-expressing cells and the highest mean expression among expressing cells (Fig. 3D). *Cxcl9* was expressed at lower levels than *Cxcl10* at this timepoint, and was predominately expressed by monocytes, DCs and neutrophils (Fig. 3D). CXCL11 is also a CXCR3 chemokine in mice, but C57BL/6 mice have a frameshift mutation rendering this gene silent (*40*).

CXCL10 and CXCL9 gradients in tissues attract immune cells such as activated T cells, B cells, NKs, and DCs in a CXCR3-dependent manner (*37, 38*). Given the enrichment of CXCR3 chemokines in the vdLN upon ChAd vaccination and the expression of CXCR3 by the activated transferred T cells (fig. S1B), we hypothesized that ChAd could improve activated CXCR3+ T-cell trafficking to the vdLN in an antigen-independent manner (Fig. 3E). Indeed, both ChAd-P1A and ChAd-i.a. immunizations increased CD8+ T cell numbers in the vdLN (Fig. 3F). We next investigated if we could identify P1A-specific T cells among the accumulated T cells and found that both ChAd-P1A and ChAd-i.a. significantly increased the number of P1A-specific T cells in the vdLN (Fig. 3G). Neither vaccine increased T-cell numbers in the non-draining LN relative to ACT alone (Fig. 3, F and G). We also investigated how the vectors with and without the antigen could affect the antigen-specific T-cell phenotypes in the LNs. The P1A-specific T cells were divided into 5 clusters, namely naïve-like, central memory, central memory CXCR3^hi^, effector CD62L^mod^ and effector CD62L^low^ (Fig. 3H and fig. S5A) according to their expression of KLRG1, CD44, CXCR3, CD62L, CD127 and PD1 (Fig. 3I and fig. S5A). Though the proportions of memory and naïve-like P1A-specific T cells in the vdLN of mice that received ChAd-i.a. were similar to those that received T cells alone (fig. S5B), the number of cells populating all clusters increased significantly in the ACT plus ChAd-i.a. compared to the ACT group alone (fig. S5C). In the ChAd-P1A vaccinated group, a proportion of cells acquired effector phenotypes with high expression of PD1, CXCR3, and CD44 and low expression of CD127 (Fig. 3, H and I and fig. S5, B and C), indicating that in addition to promoting T-cell migration to the vdLN, ChAd-P1A also promoted T-cell activation through antigen presentation.

Therefore, 24 hours after ChAd vaccination, populations of monocytes, neutrophils, DCs and γδ T cells express CXCL10, and neutrophils, monocytes and DCs express CXCL9 in the vdLN. These cytokines were associated with a higher proportion of total CD8+ T cells and antigen-specific T cells in this lymphoid tissue, indicating that ChAd can promote antigen-specific T-cell accumulation in the vdLN independent of a cognate antigen.

### T-cell homing to and subsequent egress from LNs are crucial for vaccine-mediated T cell expansion in vivo

Because ChAd vaccination leads to a local expansion/infiltration of the P1A-specific T cells in the vdLN, we investigated whether pharmacological perturbations of T-cell trafficking to/from the LNs would be sufficient to hinder vaccine-mediated T-cell expansion. We used an antibody blocking CD62L (αCD62L) to prevent homing of T cells to the lymph nodes (*41, 42*) and the small molecule FTY720 (fingolimod) to block the sphingosine-1-phosphate receptor, which is used by T cells to egress LNs (and other lymphoid tissues) along sphingosine-1-phosphate gradients (*42, 43*) (fig. S6A). While P1A-specific T cells were more abundant in the blood 6 days after ACT with ChAd-P1A compared to ACT alone, this expansion was abolished with αCD62L or with FTY720 (fig. S6B). These results indicated that homing of transferred T cells to LNs was required to allow their expansion following ChAd prime. These observations are in line with previous reports showing that combining ACT with vaccination in mice with impaired secondary lymphoid tissues failed to control tumor challenge (*3*).

After MVA-P1A boost, reductions in the number of circulating P1A-specific T cells were also observed after treatment with αCD62L or FTY720, although they were less pronounced (fig. S6C). In the vdLN, 9 days after the MVA-P1A boost, P1A-specific T cells were reduced after αCD62L, as expected, but also, surprisingly, after FTY720 (fig. S6D). Of note, FTY720 and αCD62L also reduced total blood CD8+ T cells at day 6, but on day 14 this was the case only for FTY720 (when comparing to T cells plus vaccination) (fig. S6, E to G).

These results suggest that perturbations in ACT migration to and from the LN are deleterious for antigen-specific T-cell expansion following both priming and boosting vaccination.

### CXCR3 expression promotes T-cell accumulation in lymph nodes and tumors

To confirm the link between CXCL10 and CXCL9 expression in the vdLN and the antigen-specific T-cell accumulation in response to ChAd vaccination, we generated *Cxcr3*^-/-^ mice in the OT-I background by crossing *Cxcr3^-/-^* mice with OT-I CD45.1 mice. We validated their phenotype by stimulating splenocytes with the cognate SIINFEKL peptide and IL2 (fig. S7A). As expected, CXCR3 was only detected on wt and not *Cxcr3*^-/-^ T cells (fig. S7B). A comparable proportion of T_CM_ and T_EFF_ cells was observed across the different genotypes after activation (fig. S7C), indicating that CXCR3 is not required for the cells to transition from naïve to central memory and effector phenotypes ex vivo.

Next, we investigated how CXCR3 affects T-cell distribution across tissues and how it is impacted by ChAd vaccination. To this end, we co-transferred an equal number of wt (CD45.1+) and *Cxcr3*^-/-^ OVA-specific T cells (CD45.1/2+) stained with cell-trace (CT) to melanoma-bearing mice and monitored the transferred (CD45.1+) T-cell distribution in various tissues in response to vaccination (Fig. 4A and fig. S7D). *Cxcr3*^-/-^ T cells were the predominant genotype in the blood (Fig. 4B). However, the proportion of the CXCR3-deficient T cells was dramatically reduced in the vdLN compared to the wt T cells after vaccination, indicating that CXCR3 expression favors T-cell expansion/migration to the vdLN (Fig. 4, B to D). This difference was not due to the incapacity of the *Cxcr3*^-/-^ T cells to proliferate in the vdLN, because both types of T cells showed proliferation in the vdLN in response to vaccination (Fig. 4, E and F). Moreover, CXCR3-deficiency had a small to no impact on T-cell abundance in the spleen and the inguinal (tumor draining) LN but resulted in a massive reduction in tumor infiltration even in the context of vaccination (Fig. 4, B to D).

**Fig. 4.**
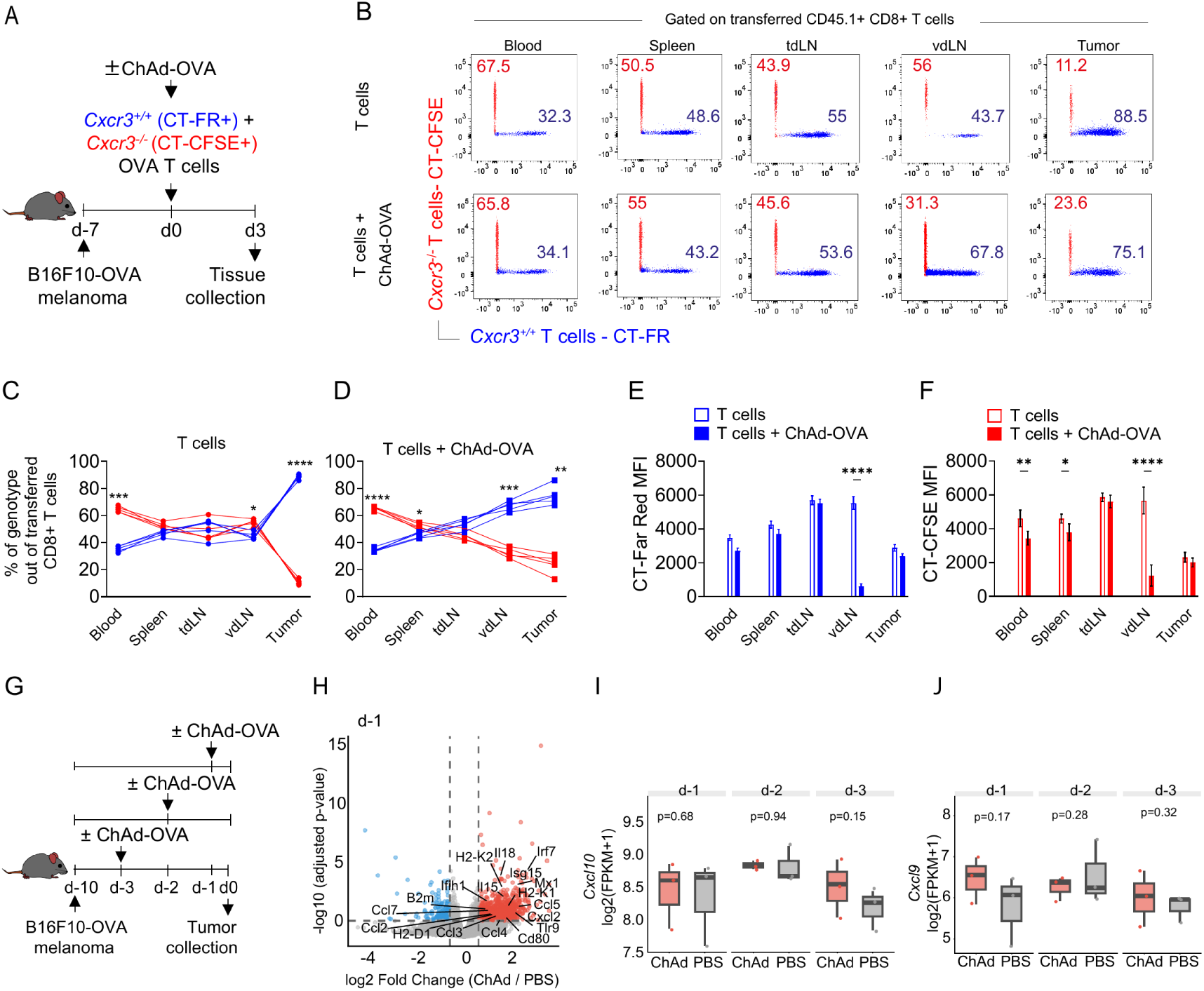
CXCR3 expression confers an advantage for T cells to accumulate in the vaccine-draining lymph node and infiltrate in tumors. **(A)** OT-I *Cxcr3*^+/+^ (CD45.1+) and *Cxcr3*^-/-^ (CD45.1/2+) T cells were stained with cell trace (CT) Far Red (FR) or CFSE, respectively, and injected in tumor-bearing mice alongside ChAd-OVA vaccination. Blood, spleen, tumor-draining LN (tdLN), vaccine-draining LN (vdLN) and tumors were collected 3 days after treatment. **(B)** Transferred T cells (CD45.1+ or CD45.1/2+) were gated out of CD8+ T cells and their respective CT staining are displayed as representative plots of *Cxcr3*^-/-^ T cells (red) or wt T cells (blue) in the indicated tissues. Numbers indicate the percentage of cell genotype out of transferred T cells. **(C-D)** Quantification of data shown in (B) in the T cell alone (C) or T cells + ChAd-OVA (D) groups. **(E-F)** CT-Far Red Median Fluorescence Intensity (MFI) of the CD45.1+ *Cxcr3*^+/+^ OT-I cells **(E)** or CT-CFSE of the CD45.1/2+ *Cxcr3*^-/-^ OT-I cells **(F)** in the respective tissues. **(G)** Mice were challenged with B16F10-OVA melanoma tumors on day -10, vaccinated on day -3 or -2 or -1, and the tumors were collected on day 0 for bulk RNA-seq. **(H)** Volcano plot displaying the differential expression between ChAd and PBS groups (day -1). DESeq2 log2 fold changes (ChAd/PBS) and Benjamini-Hochberg adjusted p-values were used for visualization. Genes with |log2FC| > log2(1.5) and padj < 0.05 were considered differentially expressed. Innate immune sensing genes (*Tlr9*, *Ifih1*), type I interferon response genes (Irf7, Mx1, Isg15), chemokines (*Cxcl2*, *Ccl2*, *Ccl3*, *Ccl4*, *Ccl5*, *Ccl7*), pro-inflammatory cytokines (*Il15*, *Il18*), and MHC-I antigen presentation and co-stimulatory genes (*H2-K1*, *H2-K2*, *H2-D1*, *B2m*, *Cd80*) are highlighted at day -1. (I) *Cxcl10* and (J) *Cxcl9* expression from the same bulk RNA-seq experiment described for Fig. 4G are displayed as boxplots. Gene expression is shown as log2(FPKM + 1). Adjusted p-values (Benjamini-Hochberg) from DESeq2 are shown.

As we observed a fast accumulation of transferred *Cxcr3*^+/+^ but not *Cxcr3*^-/-^ T cells in the tumors even in the absence of vaccination, we defined the cell populations expressing CXCL10/9 in the B16F10 TME. We reanalyzed scRNA-seq data of B16F10 tumors (*44*) and identified macrophages, monocytes, regulatory T cells (Tregs) and DCs as the clusters with the highest mean expression of *Cxcl10*, whereas macrophages and B cells had the highest *Cxcl9* expression (fig. S8A). In an independent B16F10 scRNA-seq dataset (*45*), inflammatory macrophages were also the main cells expressing *Cxcl10*/9. Interestingly, *Cxcl10* (but not *Cxcl9*) was also detected in the tumor cells (fig. S8B).

We have previously reported that after ChAd-P1A and MVA-P1A vaccination, there is upregulation of *Ifng*, *Cxcl9*, *Cxcl10*, *Ccl3*, *Ccl5* and *Xcl1* in the TME (*27*). To investigate how one dose of ChAd would affect the TME, particularly the expression of *Cxcl10/9*, we performed bulk RNA-seq of B16F10-OVA tumors 1, 2 and 3 days after ChAd i.m. (Fig. 4G). Chemokines (*Ccl5*, *Cxcl2*, *Ccl4*, *Ccl3*, *Ccl2*, *Ccl7*), cytokines (*Il18*, *Il15*), MHC-I molecules (*H2-D1*, *H2-K2*, *H2-K1*, *B2m*), co-stimulatory receptors (*Cd80*) and IFN-associated genes (Irf7, Mx1, Isg15) were upregulated only on day 1 after ChAd vaccination (Fig. 4H and fig. S8, C to E). However, consistent with the TME T-cell accumulation data, there was no significant difference in the levels of *Cxcl10* and *Cxcl9* transcripts 1-3 days after ChAd in B16F10-OVA (Fig. 4, I and J), possibly due to the already high expression of the chemokines at baseline.

Altogether, the B16F10 TME has a high baseline expression of *Cxcl10* and *Cxcl9*, which are not increased with only 1 dose of ChAd, suggesting that the vaccine instead is an important agent in promoting T cell accumulation in the vdLN through the CXCL10/9-CXCR3 axis in this model.

### CXCR3 expression is required for ACT and vaccination combination efficacy

We showed that ChAd promoted antigen-specific T cell enrichment in the vdLN via the CXCL10/9-CXCR3 axis. Furthermore, IFNα signature was significantly increased in the vdLN (Fig. 3C and fig. S4F) and tumors after ChAd (fig. S8C), and the IFNα receptor, IFNAR, was previously shown to be required for ChAd immunogenicity (*28*). Therefore, we investigated whether the anti-tumor effect of ACT in combination with ChAd/MVA was dependent on the interaction of CXCL10 or IFNα with their receptors, CXCR3 and IFNAR, respectively. Thus, we treated tumor-bearing mice with ACT and the viral vector vaccines, as previously, and administered saturating doses of either αIFNAR or αCXCR3 antibodies (Fig. 5A).

**Fig. 5.**
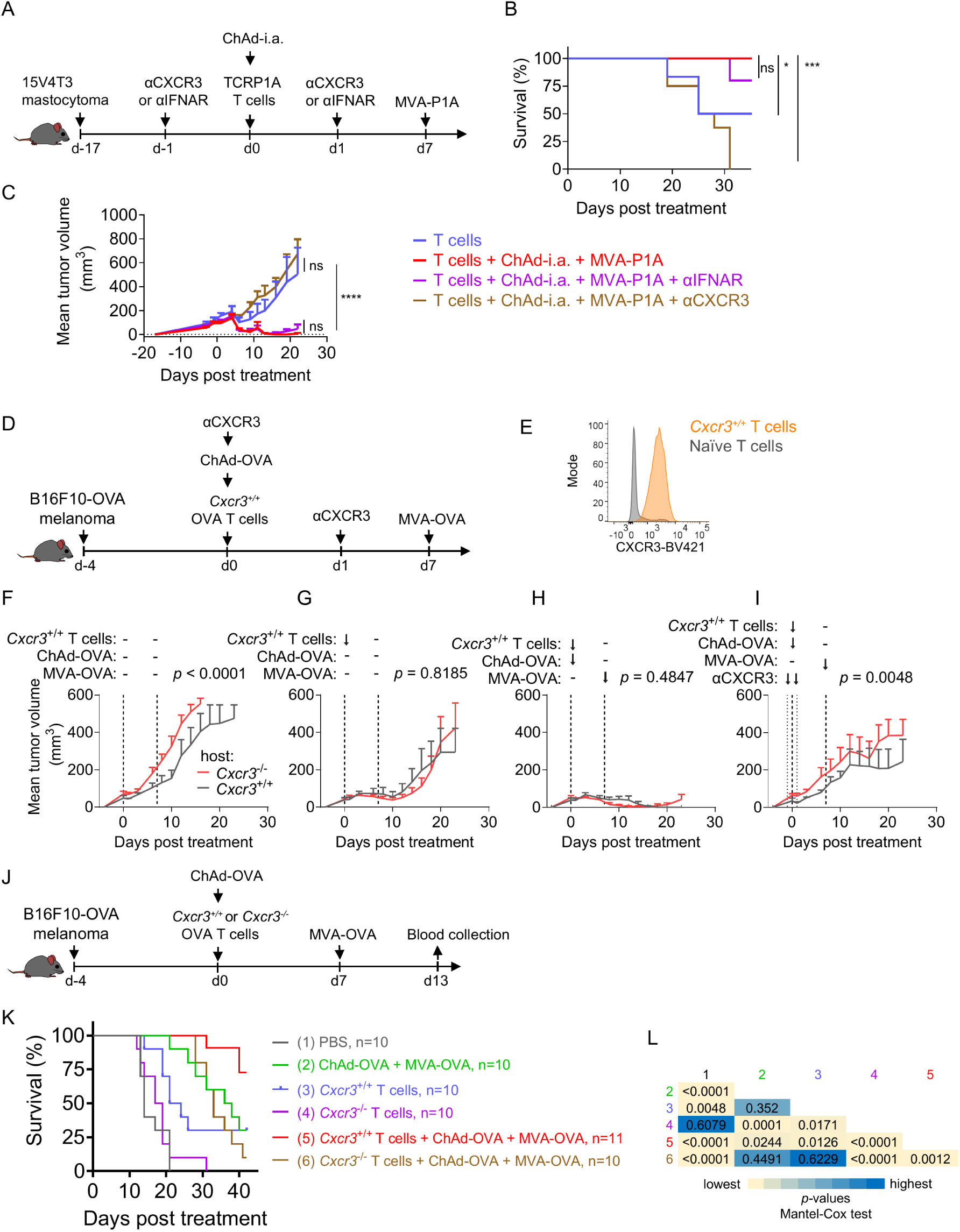
The efficacy of ACT combined with vaccination requires CXCR3 on the transferred T cells, but not host CXCR3 or IFNAR signaling. **(A)** 15V4T3 mastocytoma-bearing mice were treated with TCRP1A T cells and ChAd-i.a. and boosted with MVA-P1A, alongside saturating concentration (500 μg per timepoint) of αCXCR3 or αIFNAR antibodies (day -1 and +1). (B) Kaplan-Meier survival curve and **(C)** mean tumor growth kinetics post treatment. (**D**) B16F10-OVA tumors were implanted in CXCR3-competent or deficient mice. *Cxcr3*^+/+^ T cells were transferred alongside or not ChAd-OVA, αCXCR3 antibody. An MVA boost was given 7 days later. (**E**) Representative histogram plot of CXCR3 expression on the transferred T cells compared to naïve T cells. (F-I) Mean tumor volumes in *Cxcr3*-competent and deficient mice. (**J**) B16F10-OVA-bearing mice were treated with either *Cxcr3*^+/+^ or *Cxcr3*^-/-^ OVA-specific cells alongside ChAd-OVA and MVA-OVA. (**K**) Kaplan-Meier survival curves and (**L**) the Mantel-Cox test *p-values*. p-values below 0.05 were considered statistically significant.

The anti-tumor effect of the ACT combined with vaccines was not affected by blocking IFNAR (Fig. 5, B and C), possibly due to a compensatory mechanism in which transient IFNAR blockade increases IFNγ (and DC-derived CXCL9/10 as a result) (*39*). In contrast, αCXCR3 completely abolished the tumor control resulting from the combination therapy (Fig. 5, B and C and fig. S9, A to D), highlighting the important role of CXCR3 across tumor models. We further validated the key role of CXCR3 in the B16F10-OVA melanoma model. Furthermore, to determine whether CXCR3 expression was crucial on the transferred T cells or on the host cells for combination therapy efficacy, we implanted B16F10-OVA tumors in *Cxcr3*^+/+^ and *Cxcr3*^-/-^ mice and treated them with ACT of OT-I CXCR3+ CD8+ T cells combined with ChAd-OVA/MVA-OVA vaccination (Fig. 5, D and E). This combination controlled tumor growth equally well in CXCR3-competent and deficient mice, and in both cases this anti-tumor effect was abolished in presence of αCXCR3 (Fig. 5, F to I, and fig. S9, E to L). This result indicated that the effect of CXCR3 blockade in preventing efficacy of the combination therapy was mediated by its effects on the transferred T cells rather than on host cells.

To further validate this finding, we transferred the activated CXCR3+ or *Cxcr3*^-/-^ OVA-specific CD8+ T cells to melanoma-bearing wt mice and vaccinated them with ChAd/MVA-OVA vaccines (Fig. 5J). On day 13, peripheral blood mononuclear cells (PBMCs) were restimulated ex vivo with the cognate SIINFEKL peptide. Comparable numbers of degranulating, IFNγ-producing (CD107a+IFNγ+) transferred *Cxcr3*^-/-^ and *Cxcr3*^+/+^ T cells were detected in the blood in response to vaccination (fig. S10A). To compare their differentiation potential in an unbiased, high-dimensional manner, we visualized SIINFEKL-stimulated CD8⁺ T cells by UMAP. The transferred (CD45.1+) cells of both genotypes occupied the same effector landscape, defined by memory/differentiation (CD44, CD62L, CD127, KLRG1) and functional (IFNγ, TNF, IL2, CD107a) markers (fig. S10, B and C). Unsupervised clustering of the transferred CD45.1⁺ cells resolved the same subpopulations in both genotypes, with no significant difference in their relative frequencies between *Cxcr3*^+/+^ and *Cxcr3*^-/-^ T cells (fig. S10, D to F). Thus, *Cxcr3*^-/-^ T cells retained the capacity to expand, produce effector cytokines, and acquire effector/effector-memory states equivalent to their wt counterparts, indicating intact proliferation and functional differentiation in vivo.

As shown earlier, combination therapy of wt T cells plus vaccines resulted in superior tumor control and survival compared to any monotherapy alone (Fig. 5, K and L and fig. S10, G to K). Crucially, however, adding *Cxcr3*^-/-^ T cells to vaccination gave no benefit over vaccination alone (Fig. 5, K and L and fig. S10, K and L), confirming that CXCR3 expression on the transferred T cells is required for their anti-tumor effects in the context of vaccination. CXCR3-deficient T cells alone also had a lower capacity to control tumors compared to their wt counterparts (Fig. 5, K and L and fig. S10, I and J), consistent with previous reports (*46*).

Altogether, the anti-tumor synergy of ACT with ChAd/MVA against established, solid tumors depends on CXCR3 expression by the transferred T cells. CXCR3 favors T-cell accumulation in the vaccine-draining lymph nodes as well as tumor infiltration and is therefore the critical axis of this combination therapy.

## DISCUSSION

ACT has limited efficacy against solid tumors, in part because tumor-reactive T cells fail to undergo sufficient expansion to sustain durable anti-tumor responses. Here, we show that heterologous ChAd/MVA vaccination overcomes this limitation by promoting robust expansion of transferred T cells in lymphoid tissues, followed by their accumulation within tumors, resulting in curative anti-tumor immunity against well-established solid tumors without the need for host pre-conditioning. We and others have shown that trafficking to lymphoid tissues, such as the LNs, is a crucial step for T-cell expansion in the context of vaccination (*3*). However, even T cells capable of migrating to lymphoid tissues (due to their CD62L expression) have limited expansion in tumor-bearing hosts, suggesting that tumor-derived antigens drained to the LNs are insufficient to expand existing tumor-reactive T cells at levels that can induce sustained anti-tumor responses. Importantly, we observed that this lack of efficacy can be overcome by the administration of ChAd/MVA, which results in a robust expansion of the T cells in lymphoid tissues and subsequent tumor infiltration. Thus, ChAd/MVA vaccination emerges as a new therapy modality to be combined with ACT due to its capacity to expand cognate T-cell products and promote curative levels of anti-tumor T cells across tissues, with a long-lasting memory effect.

Once sufficient T cells have been generated, another significant obstacle for ACT is the cellular infiltration into tumors. The mastocytoma tumor model used herein has been reported as a “cold tumor” due to the poor infiltration of CD4+ and CD8+ T cells (*27*). We observed that a ChAd immunization enhanced the magnitude of inflammatory cytokines produced after MVA vaccination (e.g. IFNγ, CXCL10), thereby promoting CD8+ T cell infiltration. The heterologous prime and boost strategy favors not only the tumor-specific T cell responses, but also the cytokines that promote their function, localization and migration across tissues (*47, 48*). Importantly, our data support a model in which the vdLN acts as a critical amplification hub for adoptively transferred T cells. ChAd/MVA vaccination establishes a chemokine-rich environment that actively recruits CXCR3-expressing T cells and promotes their accumulation within the vdLN, where they can undergo further expansion and reactivation. In fact, we observed that blocking T-cell trafficking significantly reduced the expansion of antigen-specific T cells in response to priming and boosting, highlighting that T-cell migration within and between LNs is required for T-cell expansion in the context of T-cell therapies and vaccination. Our work therefore emphasizes that therapies to be combined with ACT should promote not only T-cell infiltration into tumors, but also the accumulation, restimulation, and amplification of transferred T cells in LNs.

In humans, CXCR3 expression on CD8+ T cells is predictive of improved overall survival in melanoma, breast, head and neck tumors (*49*). These results are in line with the reduced tumor control we observed when the combination of vaccines and ACT was administered with αCXCR3 or when the T cells were CXCR3-deficient. CXCR3 plays an important role in T-cell positioning in the LN and spleen. For example, CXCR3-deficient T cells have limited access to recently arrived CXCL10-producing DCs in the LN interfollicular region, negatively impacting the formation of effector T-cell phenotypes (*38, 50–52*). Indeed, naïve CXCR3-deficient OVA T cells showed reduced expansion in response to peptide stimulation in comparison to co-transferred wt counterparts in blood and LNs (*51*). Furthermore, the efficacy of αPD1 therapy relies on intact CXCR3 functionality in the TME (*50*), and patients with higher levels of the CXCR3 ligands in the plasma were more likely to respond to neoadjuvant immunotherapy against melanoma (*50, 53*), highlighting that many immunotherapies rely on the CXCL10-CXCR3 immune axis. We expanded on these results, showing that deficiency in CXCR3 signaling is also detrimental in the context of ACT and cancer vaccine therapy in tumor-bearing hosts and that CXCR3 expression on adoptively transferred T cells is necessary for their trafficking to vdLNs where they can be further expanded after vaccination.

Interestingly, while IFNAR-deficient mice have impaired antigen-specific T cell responses following ChAd vaccination (*28*), we show that IFNAR signaling shortly after vaccination is dispensable for the combinatorial efficacy of ACT and vaccination. This suggests that IFNAR is crucial for T-cell priming, but not for reactivation of memory cells in the context of vaccination. Instead, CXCR3 was the crucial factor in the combination therapy. High levels of CXCL10 induction after ChAd were reported in the serum of mice (*28*), at the transcript level in human blood after vaccination (*35*) and in human myeloid cells in vitro (*54*). The chemokine can be induced by IFNα and IFNγ (*39, 55, 56*). In our study, monocytes, neutrophils and γδ T cells were identified as the main sources of CXCL10 in the vdLN. We showed that even ChAd encoding an irrelevant antigen promoted the accumulation of transferred antigen-specific T cells in the vdLN, raising the possibility that the vector could synergize with ACT in an antigen-independent manner. Indeed, ChAd-i.a. prime combined with ACT and a cognate MVA boost was as effective as the fully cognate regimen in increasing mice survival. Although this combination was not consistent across the tumor models we investigated, understanding the mechanisms behind the failure of the ChAd-i.a. in some models could pave the way toward antigen-agnostic, off-the-shelf ChAd vaccines for immunotherapies. Consistent with this concept, mRNA-LNP-based SARS-CoV-2 vaccination has been associated with increased survival in humans even when given 100 days prior to ICI therapy (*57*), and influenza vaccination has been associated with longer survival in patients receiving ICI (*58*). This synergy was associated with pro-inflammatory mechanisms induced by vaccination independent of the antigen present in the vaccine (*57*). Whether adenovirus vaccination can favor improved outcomes for patients receiving immunotherapy still needs to be defined in retrospective cohort studies or by clinical trials.

We previously demonstrated that ChAd and MVA viral vector vaccination can treat solid tumors when combined with ICI therapy, such as αPD1 (*27*). This suggested that T-cell exhaustion might be one of the mechanisms limiting vaccine-mediated anti-tumor T-cell efficacy, and encouraged us to combine ACT therapy with αPD1. However, this combination did not enhance the anti-tumor capacity of ACT. This suggests that the PD-1/PD-L1 axis is not necessarily a crucial mechanism exploited by these tumors to escape T-cell mediated killing in the context of ACT. Crucially, we show that the combination of ACT and viral vector vaccines works efficiently in a model where αPD1 failed to enhance ACT, thus providing a promising alternative for patients who do not respond to ICI plus ACT.

While our study demonstrates robust anti-tumor effects of combining ACT with ChAd and MVA viral vector vaccines in murine models, there are limitations to consider. First, we used ACT derived from naïve splenic T-cell precursors, which may not recapitulate ACT conditions found in clinical settings. TILs used for ACT in the clinic comprise T cells that are isolated from the highly inhibitory TME and therefore may present limited stemness, with a high degree of terminal differentiation and exhaustion. Nonetheless, there are protocols in place to preserve T-cell stemness while promoting stimulation and growth. For example, T-cell stemness and central memory status can be achieved by culturing T cells in the presence of IL7 and IL15 (*13*), IL21 (*59*), or by pharmacological activation of Wnt signaling (*14*). Second, we targeted one antigen per tumor model, which may not be sufficient to eliminate highly heterogeneous tumors. However, epitope spreading, which is the generation of tumor-specific T-cell responses against antigens not directly targeted by the original therapies, has been reported following both CAR T-cell therapy and therapeutic vaccination (*18, 60*). Epitope spreading was associated with IFNγ production by the transferred T cells (*18*). Whether the enhanced persistence and intratumoral accumulation of CXCR3-expressing T cells induced by our vaccination strategy also promotes epitope spreading warrants further investigation. Our findings are particularly relevant to clinically approved TCR-T cell therapy afamitresgene autoleucel, which targets an HLA-A2-restricted MAGE-A4 peptide and is approved for patients with previously treated, unresectable or metastatic synovial sarcoma (*5*). Our data provide a rationale for evaluating afamitresgene autoleucel in combination with ChAd/MVA-MAGE-A4 vaccination in patients with MAGE-A4-expressing tumors. More broadly, combining vaccines with ACT targeting multiple antigens might also improve therapeutic efficacy by reducing immune escape. In a recent phase I trial, 17 heavily pretreated ovarian cancer patients who received vaccine-primed, ex vivo-expanded blood autologous T cells in combination with subsequent autologous tumor lysate-loaded DC vaccination had a median overall survival twice that of historical controls (*15*). This suggests that targeting multiple antigens in an ACT plus vaccines approach is a feasible and promising endeavor. Finally, while our data highlight the critical role of CXCR3 in orchestrating T-cell accumulation and localization in the context of vaccination in the LN, its role in anti-tumor immunity is likely context-dependent. For instance, elevated CXCR3 expression on Tregs in tumor-dLN and within the tumor microenvironment was shown to promote interactions with type I DCs, thereby restricting CD8+ T-cell-mediated anti-tumor responses (*61*). This suggests that the impact of CXCR3 signaling may vary depending on the immunological composition of the tumor microenvironment.

In conclusion, we demonstrate that heterologous ChAd/MVA vaccination markedly enhances TCR-T-cell therapy, resulting in durable tumor control even where PD1 blockade was ineffective. This synergy depends on CXCR3 expression by transferred T cells, which directs their accumulation in the vdLN and their infiltration of tumors. These findings identify CXCR3 as a product-intrinsic determinant of response to ACT plus vaccination and a potential biomarker of therapeutic efficacy. These findings provide a rationale for clinical testing of viral-vector vaccines, such as ChAd/MVA-MAGE-A4, in combination with TCR-T therapies (e.g., afamitresgene autoleucel), and support the selection or engineering of T cells that retain CXCR3.

## Acknowledgments

We would like to thank Dr. Didier Colau and Dr. Vincent Stroobant for generating the tetramers used in this project; Dr. Pedro Gomez for his assistance with mice shipping. We would like to thank the Functional Genomics Facility staff, past and present, for mice husbandry and maintenance: Heather Chandler, Douglas Passos, Tom B, Luke Harris, Peter Hinton, Luisa James, Julia Roberge, Laura Clarke, Alison Chandler and Viv Clark.

## List of Supplementary Materials

Materials and Methods

Figs. S1 to S10

Tables S1 and S2

References (*62–82*)

## Author contributions

Conceptualization: BVDE, CL, VPA

Methodology: BVDE, CL, VPA, RF, FA, MI, AG, LG

Investigation: VPA, SAM, AW, JMc, LN, ES, SP, YY, LG, AB, CH, LY, AA

Visualization: VPA

Funding acquisition: BVDE, PK, CL

Project administration: BVDE, CL, VPA

Supervision: BVDE, CL, RF

Writing – original draft: VPA, BVDE, CL

Writing – review & editing: VPA, BVDE, CL, RF, SAM, AW, JMc, LN, ES, SP, MI, FA, AG, LG, AB, CH, LY, AA.

## Data, code, and materials availability

All data needed to evaluate the conclusions in the paper are present in the paper or the supplementary materials. The bulk RNA-sequencing data of B16F10-OVA tumors generated in this study have been deposited in the NCBI Gene Expression Omnibus (GEO) and will be released publicly on publication; the accession number is available from the corresponding authors on request. The single-cell RNA-sequencing data of the vaccine-draining lymph node are being deposited in GEO, and the accession number will be provided before publication. Previously published datasets reanalyzed in this study are available in GEO under accessions GSE307143 (*44*), GSE121861 (*45*) and GSE304583 (*35*). All analysis code used to generate the figures, together with the annotated single-cell objects needed to reproduce the single-cell analyses, is archived at Zenodo (DOI: 10.5281/zenodo.22655495) and is also available at https://github.com/vinny258/cxcr3-act-vaccine-code. Source data for all main and supplementary figure panels are deposited at Zenodo (DOI: 10.5281/zenodo.23134889) as a multi-sheet workbook and as per-panel CSV files; access to that record is available on request and it will be released publicly on publication. All viral vectors used in this study, comprising ChAd-P1A, ChAd-OVA, MVA-P1A, MVA-OVA and their irrelevant-antigen controls, are available from the Viral Vector Core Facility, University of Oxford, under a material transfer agreement. The 15V4T3, B16F10-OVA and L1210.P1A.B7-1 cell lines and the CXCR3-deficient OT-I mouse line are available from BVDE under a material transfer agreement. Materials covered by US patent 12,180,501 are available for research use under a material transfer agreement.

