## Supplementary materials for "CXCR3 on transferred T cells enables vaccine-boosted adoptive T-cell therapy in solid tumors"

**The PDF file includes:**

Materials and Methods
Figs. S1 to S10
Tables S1 and S2
References *(62–82)*

#### Materials and Methods

##### Study Design

The objective of this study was to determine whether, and by what mechanism, chimpanzee adenovirus (ChAd) and modified vaccinia Ankara (MVA) viral-vector vaccines enhance the anti-tumor efficacy of adoptive T-cell therapy (ACT) against solid tumors. These were controlled preclinical studies in two immunocompetent mouse tumor models: P1A-expressing 15V4T3 mastocytoma in B10D2D2 mice (F1 of DBA/2 x B10.D2) and B16F10-OVA melanoma in C57BL/6 mice. The endpoints were tumor volume, measured by caliper every other day, and survival, with euthanasia at a humane endpoint (tumor volume of 1000 $mm^3$ for 15V4T3 and 500-1000 $mm^3$ for B16F10-OVA). Mice were randomized into treatment groups by tumor size before the first treatment. Treatments were not performed blinded, but tumor measurements were. Except for the experiment in Fig. 3G, for which an a priori power analysis (one-way ANOVA; $\alpha = 0.05$, power = 0.80) indicated that 8-10 mice per group were required, sample sizes were not predetermined statistically and were based on prior experience with these models. No animals were excluded. Each experiment was performed 2-7 times, and the data shown are pooled from independent experiments unless indicated as representative.

##### Mice

B10D2D2 mice were generated by crossing B10.D2 males with DBA/2 females. For the experiments, F1 male and female offspring were used at 7-16 weeks old. The TCRP1A transgenic (tg) B10.D2Rag1−/− (TCRP1A mice) colony was genotyped using the hot sodium hydroxide and Tris-HCl protocol *(62)* with ear biopsies. Briefly, ear tissue was digested in 25 mM NaOH/0.2 mM EDTA for 30 minutes at 98°C, followed by the addition of 40 mM Tris-HCl and centrifugation at 4000 rpm for 3 minutes. The resulting genomic DNA was used for a Polymerase Chain Reaction (PCR) to amplify the P1A TCR$\alpha$ (forward primer V$\alpha$8: GAATTCATGCGTCCTGGCACCTGC, reverse primer J$\alpha$10: ACCAGCAATCGAGTCCCACTTCCA, 500 bp band) and P1A TCR$\beta$ chains (forward primer V$\beta$1: GAATTCATGAGCTGCAGGCTTCTC, reverse primer J$\beta$1.2: AAAAGCCTGGTCCCCGACCGAAG, 400 bp band). PCR was performed with 0.5μM primers using GoTaq® Green Master Mix (Promega, cat. #M7123). Spleens from 7-16 weeks old female offspring were used for the experiments. Both colonies were obtained from the Prof Benoit Van den Eynde laboratory at the Université Catholique de Louvain, Belgium, under the assistance of Dr Pedro Gomez. The OT-I CD45.1 colony was generously provided by Dr. Vivian Lau and Dr. Audrey Gerard (Kennedy Institute, University of Oxford). For the OT-I colony, genotyping was performed via flow cytometry by staining peripheral blood cells. Cells were stained for 20 minutes with CD45.1-FITC (BD Pharmingen, clone A20), CD8-PerCP-Cy5.5 (Invitrogen, clone 53-6.7), CD45.2-eF450 (Invitrogen, clone 104), TCR V$\alpha$2-APC (BioLegend, clone B20.1), TCR V$\beta$5.1/5.2-PE (BioLegend, clone MR9-4), and dead cell staining-eF506 (eBioscience). Spleens from 7-16 weeks old female offspring were used for the experiments. The CXCR3$^{-/-}$ (B6.129P2-Cxcr3tm1Dgen/J, strain #005796) colony was acquired from the Jackson Laboratory (USA). For this colony, experiments were performed using 11-12-week-old F1 females. Additionally, female B6 mice, aged 7-11 weeks, were purchased from Envigo, UK. CXCR3$^{-/-}$ CD45.1/2+ OT-I mice were generated by crossing CXCR3$^{-/-}$ mice with OT-I CD45.1 mice. The genotyping was confirmed by Transnetyx and flow cytometry. All the breeding was conducted at the Functional Genomics Facility at the University of Oxford.

All mice were provided with food and water ad libitum and maintained on a 12-hour light/dark cycle. Breeding and procedures adhered to the guidelines specified by the UK Animals (Scientific Procedures) Act under the Project Licenses (PPL) PB050649E and PP7109477.

Cell lines

The 15V4T3 mastocytoma (P1A+) cell line was obtained from the laboratory of B. J. Van den Eynde (University of Oxford) and the B16F10-OVA melanoma line from the laboratory of Y. Shi (University of Oxford). The 15V4T3 cells were cultured in IMDM (Gibco) and B16F10-OVA cells in DMEM (Gibco), both supplemented with 10% fetal bovine serum (FBS, Labtech), 100 U $ml^{-1}$ penicillin, 100 ng $ml^{-1}$ streptomycin (Gibco), and 2 mM L-glutamine (Gibco). The 15V4T3 cell line was thawed in complete IMDM supplemented with 40 IU $ml^{-1}$ IL3. L1210.P1A.B7-1 cells were maintained in complete DMEM with 10% FBS, 100 U $ml^{-1}$ penicillin/streptomycin, and 2 mM L-glutamine *(27)*. All cell lines were cultured at 37°C in 5% CO2 and were regularly tested for mycoplasma using the MycoAlert Mycoplasma Detection Kit (Lonza Biologics, cat. #LT07-118).

Primary T cells

Spleens from female TCRP1A transgenic mice were harvested and processed into single-cell suspensions. After washing with 1x PBS, straining, and counting the cells, MACS isolation was performed using the Miltenyi CD8+ negative selection kit (cat. #130-104-075) to enrich the CD8+ fraction according to manufacturer's instructions. Briefly, splenocytes were diluted in MACS buffer (PBS with 2 mM EDTA and 0.5% BSA) and incubated with a biotin-antibody cocktail, followed by anti-biotin MicroBeads for 10 minutes at 4°C. The cell suspension was then passed through an LS column in a magnetic field, and the flow-through containing CD8+ cells was collected, counted, and adjusted to a concentration of 5E+5 cells $ml^{-1}$ in T-cell media. The T-cell media consisted of RPMI supplemented with 10% FBS, 100 U $ml^{-1}$ penicillin, 100 ng $ml^{-1}$ streptomycin, 2 mM L-glutamine, 1 M HEPES, 100 mM sodium pyruvate, 1% non-essential amino acids, and freshly added 50 µM 2-mercaptoethanol. Subsequently, 1 mL of this T-cell suspension was added to each well containing 1E+6 irradiated (100 Gy) L1210.P1A.B7-1 cells in T-cell media *(63)* and let grow for 4 days. OVA T cells were obtained from the spleens of female OT-I CD45.1 transgenic mice. The spleens were processed similarly to the previous protocol but without CD8 selection. The cells were adjusted to a concentration of 5E+5 splenocytes $ml^{-1}$ in T-cell media, in a total volume of 30 mL in a T75 flask. SIINFEKL peptides (10 nM) and freshly thawed IL2 (20 IU $ml^{-1}$) were added to the culture. On day 3, the cells were washed twice with PBS to remove the peptides, and the cell concentration was adjusted to 1E+6 cells $ml^{-1}$ in T-cell media with fresh IL2 (20 IU $ml^{-1}$) until the next day.

Adoptive T-cell therapy

ACT was performed on day 4 following the expansion of P1A or OVA T cells. Prior to transfer, live cells were isolated using the lymphoprep (STEMCELL #07851) protocol, following the manufacturer's instructions. Briefly, 15 mL of the cell suspension was carefully layered on top of lymphoprep and centrifuged without a break. The middle layer consisting of live T cells was collected and washed twice with PBS. P1A T cells were resuspended at 1-2E+5 in 200 µL and injected intravenously (i.v.) into the tail vein of B10D2D2 mice. OVA T cells were resuspended at 1E+6 cells in 200 µL and injected i.v. into the tail vein of C57BL/6JOlaHsd mice (B6). Before injections, the animals were placed in a heating chamber at 38°C for at least 10 minutes to dilate the tail vein. For experiments in which the T cells were stained with CellTrace Proliferation Kit

(CT), the cells were incubated at a concentration of 10E+6 cells $ml^{-1}$ with 1uM of CT (Far Red, cat. C34564 or CFSE, cat. C34554) for 20 minutes at 37°C. Then, the dye was quenched with 5x the volume of PBS 2% FBS for 5 minutes and cells were washed twice with PBS and the concentration adjusted with the injections.

Viral vector vaccination

For the P1A studies, 15V4T3 tumor-bearing mice were treated between days 17 and 19 following tumor implantation. Mice were administered 1E+7 infectious units (IU) of the chimpanzee adenovirus ChAdOx1-li-P1A (ChAd-P1A) *(27)* or ChAd-DPY irrelevant antigen (ChAd-i.a.) *(27)* intramuscularly (i.m.). Mice received a booster dose seven days after the prime treatment, consisting of 1E+6 plaque forming units (PFU) of Modified Vaccinia virus Ankara expressing P1A (MVA-P1A) or not (MVA-i.a.) *(27)* i.m., unless otherwise specified. Both vectors contained the insert for the complete P1A gene. The irrelevant antigen used in both ChAd and MVA vaccines was the human DPY protein. For the OVA studies, mice received 5E+5 IU of ChAdOx1-tPA-OVA (ChAd-OVA) or ChAd-i.a. i.m. Seven days later, mice were boosted with 1E+6 PFU of MVA-tPA-OVA (MVA-OVA). Control mice received 50 µL PBS i.m. one week apart. Both vectors had the complete OVA gene sequence. The ChAd-OVA and MVA-OVA vectors were produced by the Viral Vector Core Facility, University of Oxford. Non-treated control mice received 50 µL PBS i.m. one week apart.

Tumor implantation and measurements

1E+6 15V4T3 cells were implanted subcutaneously in F1 male and female B10D2D2 mice using non-supplemented IMDM, while 5E+5 B16F10-OVA cells were implanted subcutaneously in female B6 mice using non-supplemented DMEM. The tumor growth was monitored with a caliper every other day, and the tumor volume was calculated as V = ((length (mm) x $width^2$ (mm)) x 0.52). Prior to the first treatment, the mice were randomized. After reaching 1000 $mm^3$ (15V4T3) or 500-1000 $mm^3$ (B16F10-OVA), mice were euthanized. For some experiments, the tumors were surgically removed, weighed and processed for flow cytometry analysis *(27)*.

In vivo αPD1, αCD62L, αCXCR3, αIFNAR and FTY720 administration

For αPD1 treatment, 100 µg per mouse of *α*PD1 (BioXcell, clone RMP1-14) was injected intraperitoneally (i.p.) on days 3, 6, and 10 following ACT. For in vivo αCD62L neutralization, 200 µg of *α*CD62L (BioLegend Ultra-LEAF™ Purified anti-mouse CD62L antibody, clone MEL-14, cat. #104460) was administered intravenously (i.v.) via the tail vein 2 hours prior to the T cell injection. For αCXCR3 (BioXcell InVivoMAb Armenian Hamster anti-mouse CXCR3 (CD183), clone CXCR3-173, cat. #BE0249 or BioLegend Ultra-LEAF™ Purified anti-mouse antibody, clone CXCR3-173) and αIFNAR (Ultra-LEAF™ Purified anti-mouse IFNAR-1 Antibody, clone MAR1-5A3), antibodies were diluted in PBS to a final concentration of 500 µg in 200 µL PBS and injected i.p. one day before and one day after ACT, unless otherwise specified. For BioXcell αCXCR3, the isotype control was Armenian hamster IgG (anti-glutathione S-transferase, clone PIP, cat. #BE0260), and for BioLegend αCXCR3, the isotype control was Ultra-LEAF™ Purified Armenian Hamster IgG (clone HTK888, cat. #400960). For FTY720 (fingolimod hydrochloride), 50 mg $ml^{-1}$ of stock solution in DMSO was diluted in a mixture of 10% DMSO, 40% PEG300, 5% Tween-80, and 45% PBS. A dose of 20 µg FTY720 per 100 µL per mouse was administered i.p. on days 0, 2, 4, and 6, following MedChemExpress guidelines.

Tissue processing for flow cytometry

Blood was drawn from the tail vein after incubating the mouse in a heating chamber at 38°C. Erythrocytes were removed by two cycles of 1xACK lysis (0.15 M $NH_4Cl$, 1 mM $KHCO_3$ and 0.1mM Tetrasodium EDTA) followed by 1xPBS wash *(27)*. The spleens were strained and washed with 1xPBS. After, they were incubated with 1xACK lysis buffer and washed with 1xPBS *(27)*. Popliteal and inguinal lymph nodes from the side of vaccination (draining lymph nodes, dLN) or the contralateral side were collected, smashed using glass slides into a 20 cm non-treated dish or directly into a 50mL tube, and then washed with 1x PBS. Tumors were surgically removed, weighed, and stored in RPMI. For single-cell tumor dissociation, the tumors were placed in gentleMACS C-tubes containing tumor dissociation buffer (RPMI with 20 µg $ml^{-1}$ DNase I and 175 U $ml^{-1}$ collagenase type I). The tissue was dissociated using a gentleMACS dissociator (Miltenyi Biotec, program m_impTumor_01) and incubated at 37°C for 15 minutes at 150 rpm. The resulting cell suspension was washed and strained to obtain single cells *(27)*.

Peptide stimulation and degranulation assay

Single-cell suspensions were incubated for 5 hours in complete RPMI media with 4 µg $ml^{-1}$ of either 54 15-mer overlapping peptides spanning the entire P1A protein or selected immunodominant OVA peptides (SIINFEKL and ISQAVHAAHAEINEAGR). The media also contained 2 µg $ml^{-1}$ of *α*CD28 monoclonal antibodies (eBioscience, clone 37.51, cat. #16-0281-85), 20 µg $ml^{-1}$ Deoxyribonuclease I (DNase I) (Roche, cat. #10-104-159-001), and 1:300 dilution of CD107a-PE-Cy7 (BD Pharmingen, clone #1D4B, cat. #560647). In the final 4 hours of incubation, 5 µg $ml^{-1}$ of Brefeldin A (Abcam, cat. #ab193369) was added to facilitate intracellular cytokine accumulation *(27)*.

Flow cytometry

LPYLGWLVF-H2-$L^d$-tetramer-PE (P1A(35-43)-H2-Ld-tetramer) (10 uM) and SIINFEKL-H2-$K^b$-tetramer-PE ($OVA_{(257-264)}$-H2-$K^b$) (10 µM) were produced in de Duve Institute in Belgium by Dr. Didier Colau and Dr. Vincent Stroobant. Single cells were stained as described previously *(27)*. In brief, cells were incubated with αCD16/CD32 (Bioscience, 2.4G2), washed and stained for 30 minutes to 1 hour with the relevant tetramer. Cells were washed and stained for 20 to 30 minutes with surface staining monoclonal antibodies in the presence of the viability dye (Table S1). For the staining of transcription factors, the eBioscience™ Foxp3 / Transcription Factor Staining Buffer Set (#00-5523-00) was used according to the manufacturer's protocol. In brief, cells were fixed for 30 minutes and washed once with the FOXP3 kit permeabilization buffer. Intracellular antibody cocktail was added in a volume of 50 uL and incubated overnight (ON). The cells were posteriorly washed twice with the 1x permeabilization buffer. For intracellular staining of cytokines, the BD Cytofix/Cytoperm™ Fixation/Permeabilization Kit (#554714) was used according to the manufacturer's protocol. Briefly, after surface staining, cells were washed twice in PBS and incubated with Cytofix/Cytoperm for 20 minutes. After which cells were washed twice with 1x Permeabilization wash diluted in deionized $H_2O$ ($dH_2O$). Intracellular antibody master mix was added to the cells in 50 uL and incubated for 30 minutes or ON. Then, cells were washed with the 1x Permeabilization wash twice prior to acquisition in the flow cytometer. Stained cells were analyzed using a Fortessa X20 or Cytek Aurora.

Cytokine detection in serum

For serum separation, blood was collected from the tail vein into BD Microtainer tubes (BD Vacutainer, cat. #365979) and allowed to clot at room temperature for 30 minutes. The tubes were then centrifuged at 1,300 x g for 10 minutes, and the top layer (serum) was transferred to a new tube and stored at -80°C. Serum cytokine detection was performed using the LEGENDplex™ Mouse Anti-Virus Response Panel (13-plex) with a V-bottom Plate (BioLegend, cat. #740622) according to the manufacturer's instructions. Briefly, serum samples were thawed and centrifuged to remove particulates. 25 µL of each sample was added to a well containing 25 µL of Matrix A. To each well, 25 µL of pre-mixed beads for cytokine detection were added, and the plate was incubated at room temperature for 2 hours on a shaker. After incubation, the plate was centrifuged, and the supernatant was carefully discarded. Next, 25 µL of biotin-tagged detection antibodies were added, and the plate was incubated for 1 hour with shaking. Streptavidin-phycoerythrin (SA-PE) was then added and incubated for an additional 30 minutes. After washing, the plate was run on the Fortessa X20 flow cytometer for cytokine quantification. This panel allows simultaneous analysis of IFN$\gamma$, CXCL1, TNF, CCL2, IL12p70, CCL5, IL-1$\beta$, CXCL10, GM-CSF, IL10, IFN$\beta$, IFN$\alpha$, IL6. Results were analyzed using Qognit online software.

Bulk RNA sequencing

Tissues were snapped frozen freshly with liquid nitrogen, then transferred to -80°C prior to shipping to BMKGene, which performed the RNA extraction, library preparation and sequencing. Differential expression of tumor bulk RNA-seq was assessed with DESeq2 *(64)*, and published human peripheral blood bulk RNA-seq data following ChAdOx1 vaccination *(35)* were reanalyzed with edgeR *(65)*; analysis parameters are detailed in the corresponding figure legends.

Single cell RNA sequencing

Mouse immunization and lymph node processing

Female C57BL/6 mice were purchased from Charles River Laboratories at 9 weeks of age and immunized at 10 weeks of age. Right inguinal lymph nodes, ipsilateral to the vaccination site, were collected from naïve mice (day 0) and 24 h after (day 1) intramuscular vaccination in the right quadriceps with ChAdOx1 nCoV-19 *(66)* (AZD1222; 5E+9 viral particles per mouse; $n = 5$ mice per group). Samples from day 0 and day 1 mice were processed independently. Lymph nodes were mechanically dissociated through 70 µm cell strainers using the plunger of a 5 ml syringe into complete RPMI medium comprising RPMI-1640 supplemented with 10% fetal calf serum, 1% penicillin-streptomycin, 1× non-essential amino acids, 1 mM sodium pyruvate, 10 mM HEPES, and 55 µM β-mercaptoethanol. Cell suspensions were centrifuged at 524 × $g$ for 5 min at 4°C, washed once with 10 ml complete RPMI medium, and resuspended in complete RPMI medium before transfer to a single well of a 96-well U-bottom plate.

Cell staining and sorting

Cells were washed once with FACS buffer (PBS supplemented with 0.5% BSA and 1 mM EDTA) and incubated with TruStain FcX PLUS anti-mouse CD16/32 antibody (BioLegend, clone S17011E; 0.5 μg per well, 20 μg $ml^{-1}$) for 10 min at 4°C. TotalSeq-C hashtag antibody (Biolegend; one hashtag per mouse; 0.3 μg per well, 3 μg $ml^{-1}$) was then added and incubated for 10 min at 4°C, followed by addition of the surface antibody cocktail (Table S2) to achieve a final hashtag antibody concentration of 0.15 μg per well (1.5 μg $ml^{-1}$). Cells were incubated for a further 30 min at 4°C, washed twice with FACS buffer, resuspended in sort buffer (PBS containing 1% BSA and 25 mM HEPES), and stained with 5 nM SYTOX Green Nucleic Acid Stain (Thermo Fisher Scientific) for 20 min at 4°C. Live $CD45^+$ cells were sorted on a BD FACSAria III (BD Biosciences) equipped with an 85 μm nozzle, with naïve T cells ($CD3^+CD62L^+CD44^-$) and naïve B cells ($B220^+IgD^+$) selectively downsampled during sorting to reduce their proportion in the final dataset. Sorted cells were collected into HBSS supplemented with 50% fetal calf serum and 25 mM HEPES.

Single-cell library generation and sequencing

For each timepoint, hashtag-labelled cells from five mice were pooled, centrifuged at 400 × *g* for 5 min at 4°C, and resuspended in PBS containing 0.04% BSA. Cell suspensions were loaded across four channels of a Chromium Next GEM Chip K (10x Genomics). Single-cell gene expression and Feature Barcode (hashtag) libraries were generated using the Chromium Next GEM Single Cell 5' Reagent Kits v2 (10x Genomics; CG000330, Rev. F) according to the manufacturer's instructions. Libraries were sequenced by Novogene on an Illumina NovaSeq 6000 (150 bp paired-end reads). Gene expression libraries were sequenced to a mean depth of >50,000 reads per cell.

Data analysis

Sequencing reads were aligned to the mm10 transcriptome (10x Genomics reference 2020-A transcriptome reference) and quantified using Cell Ranger count (v7.1.0) *(67)*. Downstream analyses were performed in R (v4.3.3–4.5.3) using Seurat (v5.1.0–5.4.0) *(68)* unless otherwise stated. Hashtag demultiplexing was performed using cellhashR (v1.0.3) *(69)*, with cell assignments determined from the consensus call across six methods: $BFF_{cluster}$, $BFF_{raw}$, GMM-Demux *(70)*, MULTI-seq *(71)*, Seurat HTODemux *(72)*, and DropletUtils hashedDrops *(73)*.

Cells with unique molecular identifier (UMI) counts or gene counts below the median − 2 × median absolute deviations (MAD), or mitochondrial transcript content above the median + 2 × MAD, were removed. Data from all Chromium channels were merged and normalized using sctransform (v0.4.1) *(74)*. Principal component analysis (PCA) was performed using the top 3,000 variable genes, followed by batch correction with Harmony (v1.0) *(75)* using 30 principal components and timepoint (day 0 or day 1) as the integration variable. Cells were clustered using Seurat's graph-based clustering approach (resolution 0.5). Cluster marker genes were identified using the Seurat FindAllMarkers function with the Wilcoxon rank-sum test. Cell identities were assigned using SingleR (v2.4.0) *(76)* with the ImmGen reference dataset *(77)*, followed by manual curation based on canonical marker genes.

For differential gene expression analysis across all cell types combined (Fig. 3C), raw counts were aggregated at the mouse level within each timepoint to generate pseudobulk expression profiles. Genes detected in at least 5% of cells in one or more timepoints were retained, and differential expression between day 1 and day 0 samples was assessed using DESeq2 (v1.50.2) *(64)* with the design formula ~ day. Genes with an adjusted *p-value* < 0.05 and an absolute log2 fold change > 0.5 were considered significantly differentially expressed. Gene set enrichment analysis was performed using fgsea (v1.36.2) *(78)* against Hallmark *(79)* and Reactome *(80)* gene sets retrieved from MSigDB *(81)* via msigdbr (v26.1.0) (fig. S4F). Expression of *Cxcl9* and *Cxcl10* was quantified within annotated cell populations (Fig. 3D) as the percentage of cells with detectable expression (breadth) and the mean log-normalized expression among expressing cells (intensity).

Publicly available B16F10 tumor scRNA-seq datasets (GSE307143 and GSE121861) were reanalyzed using Seurat. Detailed analysis parameters are provided in the corresponding figure legends.

Software and statistical analysis

FlowJo version 10.8 was used to analyze the flow cytometry data and generate histograms. Clustering analysis was performed using OMIQ software. High-dimensional analyses of flow cytometry data (UMAP and k-means clustering of arcsinh-transformed marker expression) were additionally performed in R, as detailed in the corresponding figure legends. Statistical analysis and data visualization were conducted in GraphPad Prism versions 10.1 to 11.0. The specific statistical methods used are detailed in the legends of each figure, and a p-value of less than 0.05 (95% confidence interval) was considered statistically significant.

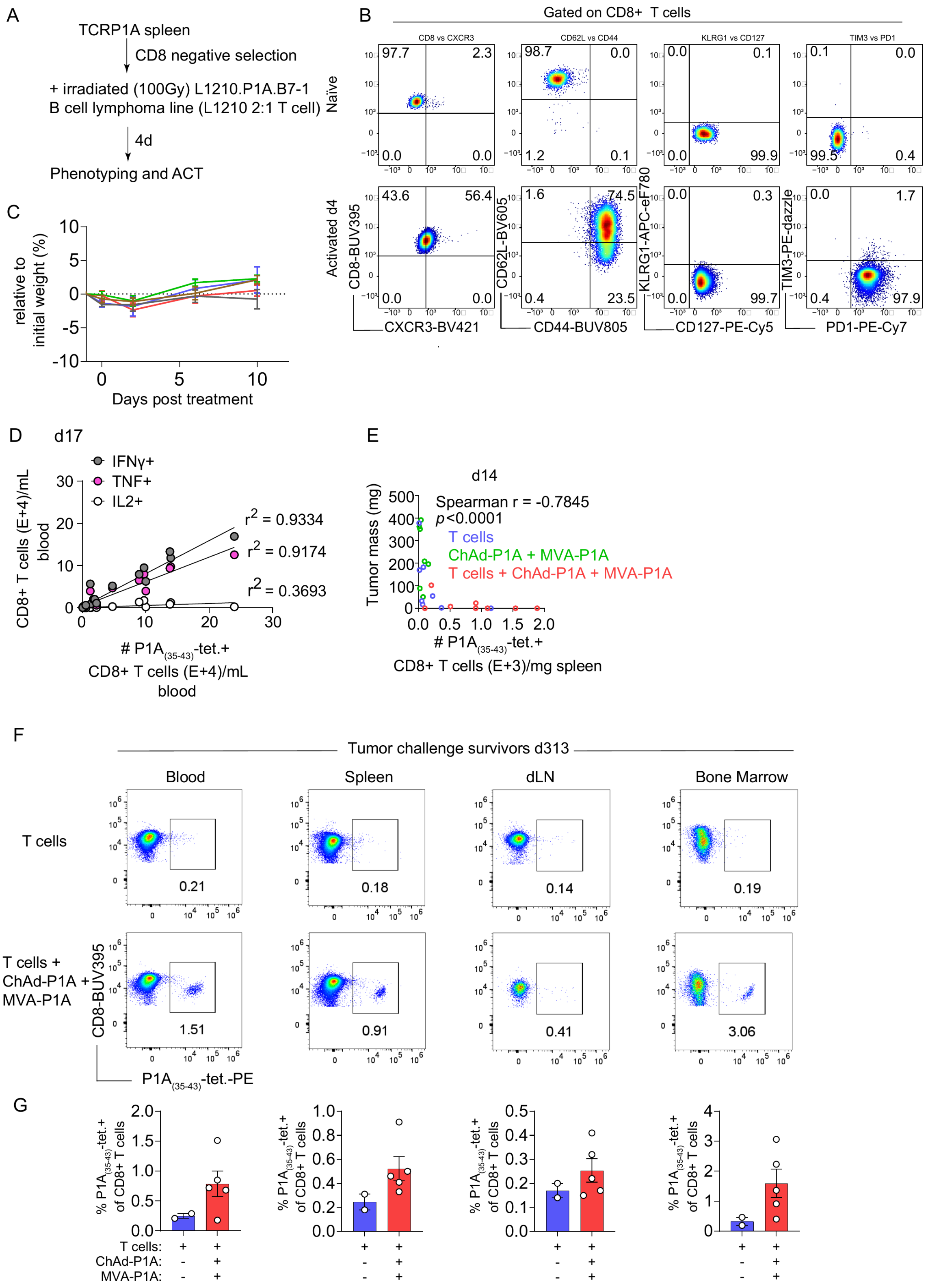

A
TCRP1A spleen
CD8 negative selection
+ irradiated (100Gy) L1210.P1A.B7-1
B cell lymphoma line (L1210 2:1 T cell)
4d
Phenotyping and ACT
B
Gated on CD8+ T cells
Naive
Activated d4
CD8-BUV395
CXCR3-BV421
CD62L-BV605
CD44-BUV805
KLRG1-APC-eF780
CD127-PE-Cy5
TIM3-PE-dazzle
PD1-PE-Cy7
C
relative to initial weight (%)
Days post treatment
D
d17
IFNγ+
TNF+
IL2+
CD8+ T cells (E+4)/mL blood
r2 = 0.9334
r2 = 0.9174
r2 = 0.3693
### P1A(35-43)-tet.+ CD8+ T cells (E+4)/mL blood
E
d14
Tumor mass (mg)
Spearman r = -0.7845
p<0.0001
T cells
ChAd-P1A + MVA-P1A
T cells + ChAd-P1A + MVA-P1A
### P1A(35-43)-tet.+ CD8+ T cells (E+3)/mg spleen
F
Tumor challenge survivors d313
Blood
Spleen
dLN
Bone Marrow
T cells
T cells + ChAd-P1A + MVA-P1A
CD8-BUV395
P1A(35-43)-tet.-PE
G
% P1A(35-43)-tet.+ of CD8+ T cells
T cells:
ChAd-P1A:
MVA-P1A:

**Fig. S1. Phenotype, anti-tumor correlation and long-term persistence of P1A-specific T cells following ACT and ChAd/MVA vaccination.** (A) T cell expansion scheme. (B) Representative plots of naïve (top row) or injected activated (bottom row, day 4) P1A T cells. (C) Weights were monitored and are shown as relative percentage with respect to weights prior to therapy. (D) Correlation between P1A-specific CD8+ T cells and cytokine production. Intracellular cytokine staining was performed after $P1A_{(35\text{-}43)}$ peptide stimulation. Coefficients of determination ($r^2$) from simple linear regression are shown. (E) Spearman correlation between tumor mass (mg) and number of $P1A_{(35\text{-}43)}$-tetramer+ CD8+ T cells per mg of spleen on day 14. (F) Representative dot plots of $P1A_{(35\text{-}43)}$-tetramer+ T cells in the respective tissues 313 days (10.29 months) after treatment in surviving mice that received T cells or T cells + ChAd/MVA vaccines. (G) Bar graphs are displaying the percentage of P1A-tetramer+ T cells out of CD8+ T cells in the indicated tissues (dLN refers to both inguinal and popliteal lymph nodes on the vaccination/tumor side).

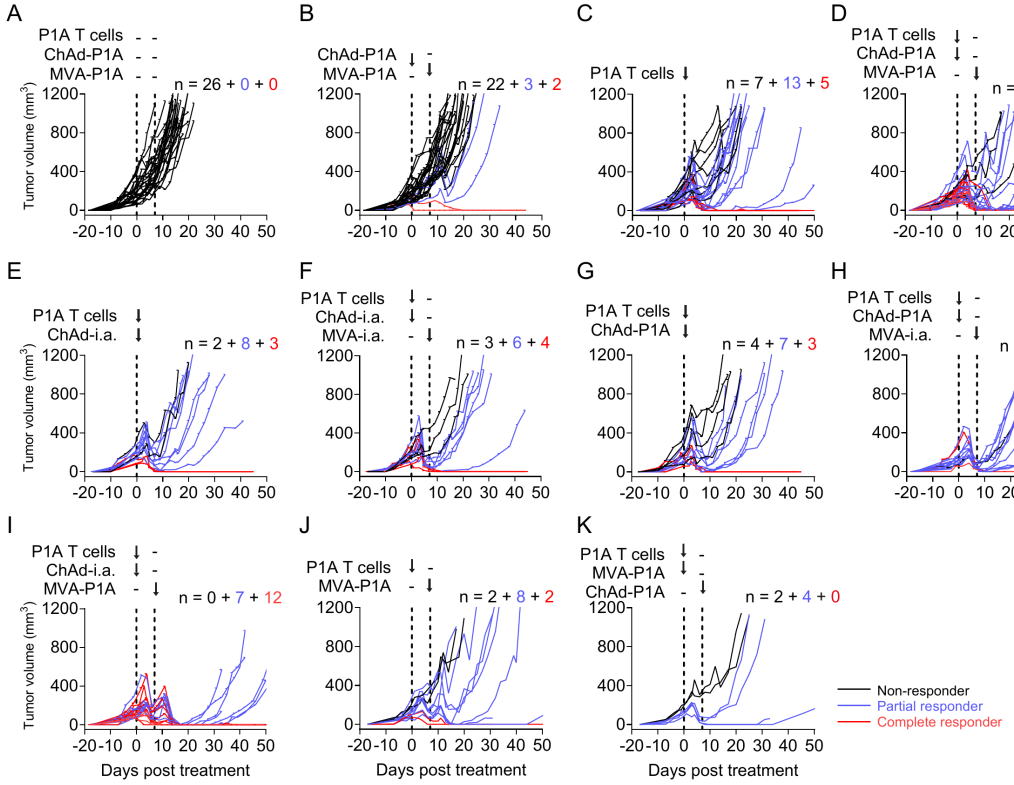
A
B
C
D
E
F
G
H
I
J
K
P1A T cells
ChAd-P1A
MVA-P1A
ChAd-i.a.
MVA-i.a.
n = 26 + 0 + 0
n = 22 + 3 + 2
n = 7 + 13 + 5
n = 3 + 14 + 15
n = 2 + 8 + 3
n = 3 + 6 + 4
n = 4 + 7 + 3
n = 1 + 10 + 2
n = 0 + 7 + 12
n = 2 + 8 + 2
n = 2 + 4 + 0
Tumor volume (mm³)
Days post treatment
Non-responder
Partial responder
Complete responder

**Fig. S2. Individual tumor growth curves of mastocytoma-bearing mice across all treatment groups. (A-K)** Plotted tumor volumes for each treatment group across the duration of the experiment. Non-responders, partial responders and complete responders are presented as black, blue or red lines, respectively.

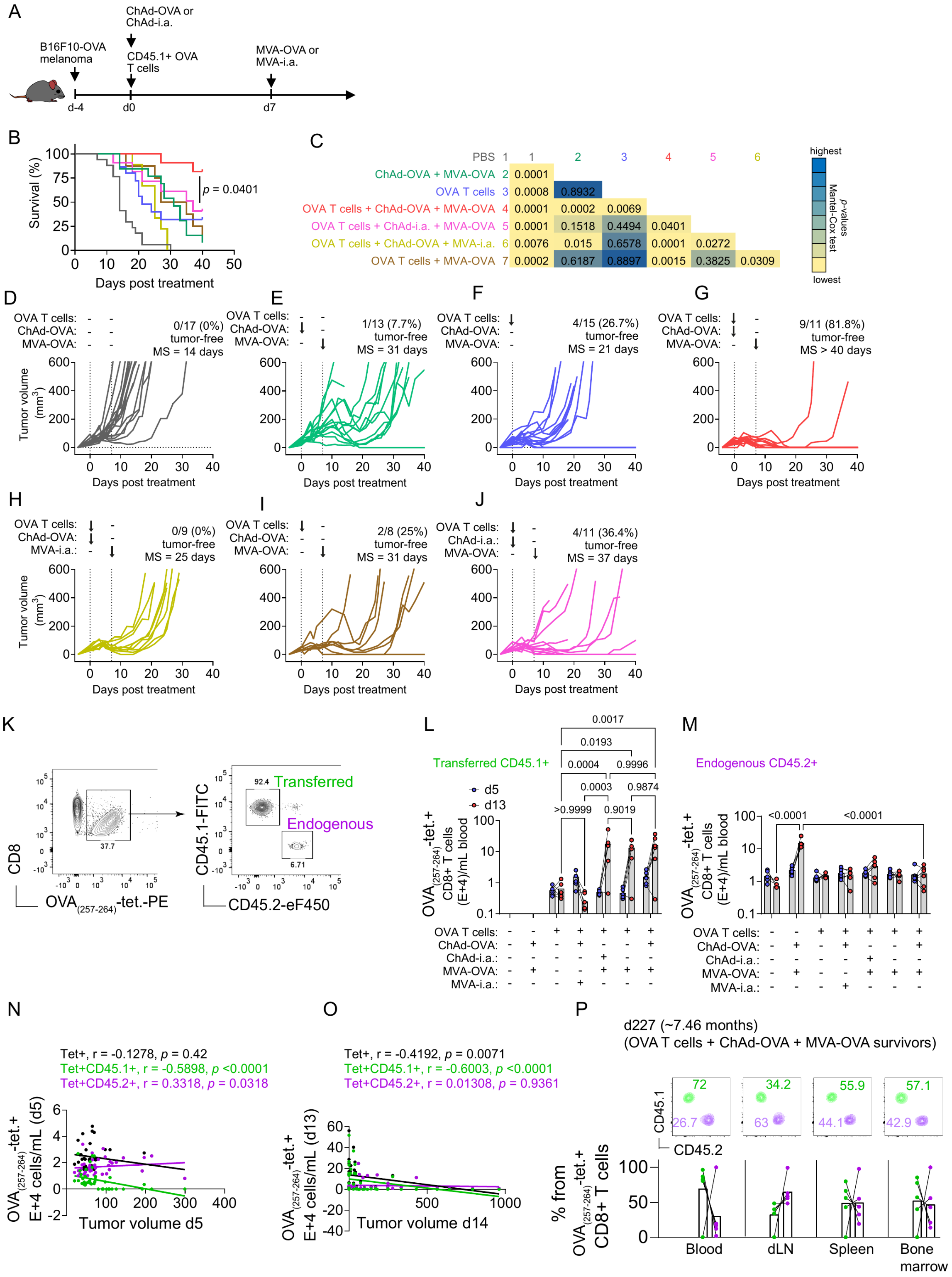

A
B16F10-OVA melanoma
ChAd-OVA or ChAd-i.a.
CD45.1+ OVA T cells
MVA-OVA or MVA-i.a.
d-4
d0
d7
B
Survival (%)
Days post treatment
p = 0.0401
C
PBS 1
ChAd-OVA + MVA-OVA 2
OVA T cells 3
OVA T cells + ChAd-OVA + MVA-OVA 4
OVA T cells + ChAd-i.a. + MVA-OVA 5
OVA T cells + ChAd-OVA + MVA-i.a. 6
OVA T cells + MVA-OVA 7
0.0001
0.0008 0.8932
0.0001 0.0002 0.0069
0.0001 0.1518 0.4494 0.0401
0.0076 0.015 0.6578 0.0001 0.0272
0.0002 0.6187 0.8897 0.0015 0.3825 0.0309
highest
p-values Mantel-Cox test
lowest
D
OVA T cells:
ChAd-OVA:
MVA-OVA:
0/17 (0%) tumor-free
MS = 14 days
Tumor volume (mm3)
Days post treatment
E
1/13 (7.7%) tumor-free
MS = 31 days
F
4/15 (26.7%) tumor-free
MS = 21 days
G
9/11 (81.8%) tumor-free
MS > 40 days
H
MVA-i.a.:
0/9 (0%) tumor-free
MS = 25 days
I
2/8 (25%) tumor-free
MS = 31 days
J
ChAd-i.a.:
4/11 (36.4%) tumor-free
MS = 37 days
K
CD8
OVA(257-264)-tet.-PE
37.7
CD45.1-FITC
CD45.2-eF450
92.4
6.71
Transferred
Endogenous
L
Transferred CD45.1+
d5
d13
OVA(257-264)-tet.+ CD8+ T cells (E+4)/mL blood
0.0017
0.0193
0.0004
0.9996
0.0003
0.9874
>0.9999
0.9019
OVA T cells:
ChAd-OVA:
ChAd-i.a.:
MVA-OVA:
MVA-i.a.:
M
Endogenous CD45.2+
<0.0001
<0.0001
N
Tet+, r = -0.1278, p = 0.42
Tet+CD45.1+, r = -0.5898, p <0.0001
Tet+CD45.2+, r = 0.3318, p = 0.0318
OVA(257-264)-tet.+ E+4 cells/mL (d5)
Tumor volume d5
O
Tet+, r = -0.4192, p = 0.0071
Tet+CD45.1+, r = -0.6003, p <0.0001
Tet+CD45.2+, r = 0.01308, p = 0.9361
OVA(257-264)-tet.+ E+4 cells/mL (d13)
Tumor volume d14
P
d227 (~7.46 months)
(OVA T cells + ChAd-OVA + MVA-OVA survivors)
CD45.1
CD45.2
72
26.7
34.2
63
55.9
44.1
57.1
42.9
% from OVA(257-264)-tet.+ CD8+ T cells
Blood
dLN
Spleen
Bone marrow

**Fig. S3. The combination immunotherapy of OVA-specific T cells with ChAd-OVA prime and MVA-OVA boost results in complete and durable tumor regression against the B16F10-OVA melanoma.** (**A**) B16 melanoma expressing the OVA antigen (B16F10-OVA) was injected s.c. on the right flank of C57BL/6 mice. Four days later, 5E+5 IU ChAd-OVA or i.a. were given i.m. and 1E+6 OVA T cells (CD45.1+) i.v. at day 0. Seven days later, mice received 1E+6 PFU MVA-OVA or MVA-i.a. i.m. (**B**) Kaplan-Meier survival curve after combination therapies and monotherapy controls. Mice survival was monitored for 40 days, and a humane endpoint was reached when tumors were around 500 $mm^3$. (**C**) Mantel-Cox test *p*-value*s* after comparison of the indicated groups. (**D-J**) Individual tumor growth curves over time. The top right side of each individual tumor growth curve shows the percentage of mice that were tumor-free by day 40 and the median survival (MS) of that group. In (**G**), the MS was undefined. (**K**) Representative plot of H-2$K^b$-SIINFEKL-tetramer+ CD8+ T cells, which were either transferred (CD45.1+) or endogenous (CD45.2+). Bar graphs show the number of transferred **(L)** or endogenous **(M)** SIINFEKL-H-2$K^b$ tetramer+ (OVA-specific) CD8+ T cells in the blood on day 5 (blue) and 13 (red). **(N-O) Spearman correlation between tumor volume** ($mm^3$) and number of total (black), transferred (green), or endogenous (purple) SIINFEKL-H-2$K^b$-tetramer+ (OVA-specific) CD8+ T cells in the blood on day 5 (**N**) or 13 (**O**). **(P)** Representative plots of transferred (green) or endogenous (purple) OVA-specific T cells in the blood, dLN, spleen and bone marrow 7.46 months after treatment. Abbreviations: IU – infectious units; PFU – plaque forming units; ChAd – chimpanzee adenovirus ChAdOx1; MVA – Modified Vaccinia Ankara virus; OVA – chicken ovalbumin; i.a. – irrelevant antigen; s.c. – subcutaneous; i.m. – intramuscular; i.v. - intravenously. *p*-values <0.05 were considered significant.

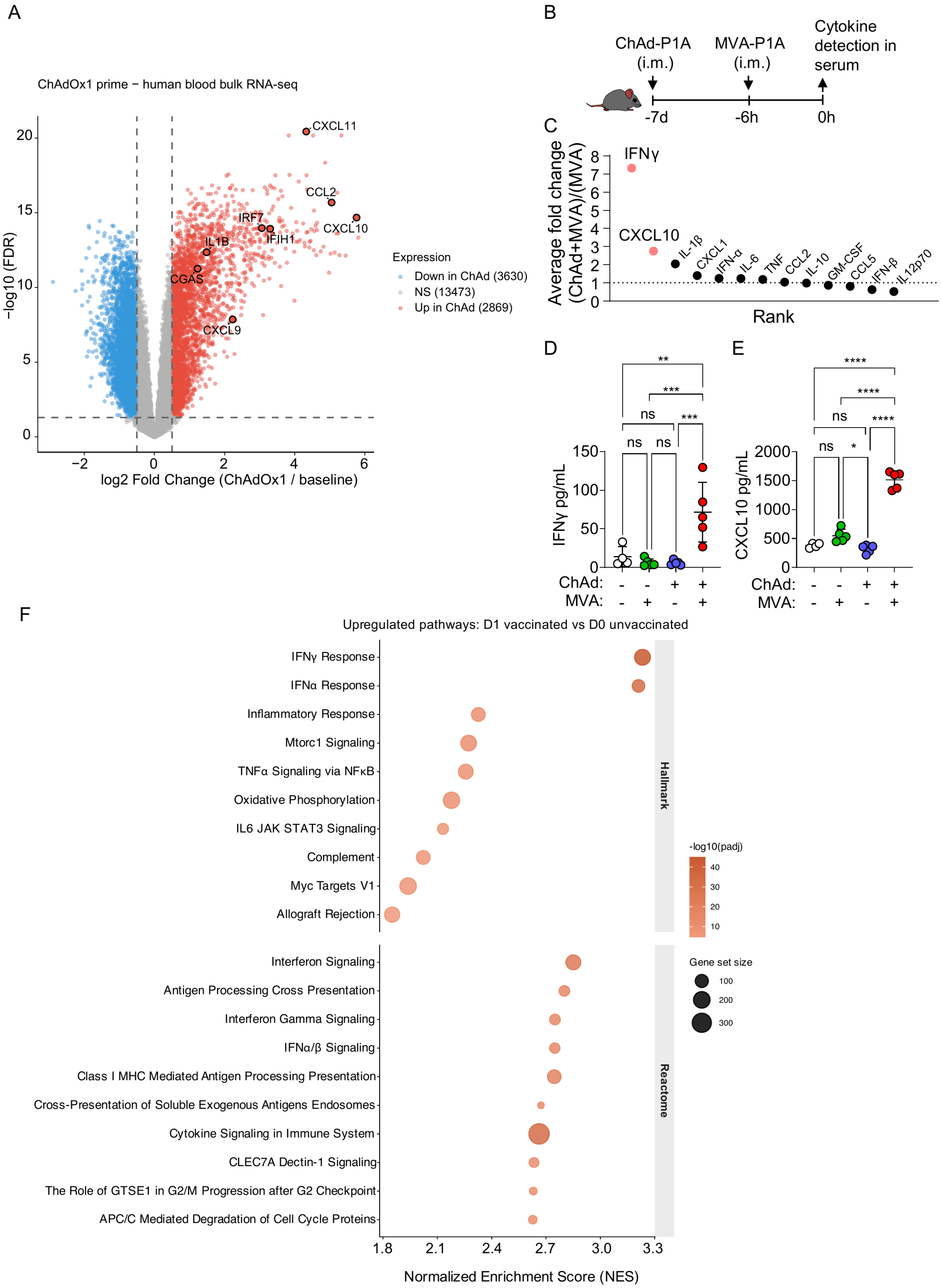
A
ChAdOx1 prime − human blood bulk RNA-seq
−log10 (FDR)
log2 Fold Change (ChAdOx1 / baseline)
CXCL11
CCL2
CXCL10
IRF7
IFIH1
IL1B
CGAS
CXCL9
Expression
Down in ChAd (3630)
NS (13473)
Up in ChAd (2869)
B
ChAd-P1A (i.m.)
MVA-P1A (i.m.)
Cytokine detection in serum
-7d
-6h
0h
C
Average fold change (ChAd+MVA)/(MVA)
IFNγ
CXCL10
IL-1β
CXCL1
IFN-α
IL-6
TNF
CCL2
IL-10
GM-CSF
CCL5
IFN-β
IL12p70
Rank
D
IFNγ pg/mL
ChAd:
MVA:
E
CXCL10 pg/mL
ChAd:
MVA:
F
Upregulated pathways: D1 vaccinated vs D0 unvaccinated
IFNγ Response
IFNα Response
Inflammatory Response
Mtorc1 Signaling
TNFα Signaling via NFκB
Oxidative Phosphorylation
IL6 JAK STAT3 Signaling
Complement
Myc Targets V1
Allograft Rejection
Hallmark
Interferon Signaling
Antigen Processing Cross Presentation
Interferon Gamma Signaling
IFNα/β Signaling
Class I MHC Mediated Antigen Processing Presentation
Cross-Presentation of Soluble Exogenous Antigens Endosomes
Cytokine Signaling in Immune System
CLEC7A Dectin-1 Signaling
The Role of GTSE1 in G2/M Progression after G2 Checkpoint
APC/C Mediated Degradation of Cell Cycle Proteins
Reactome
-log10(padj)
Gene set size
Normalized Enrichment Score (NES)

**Fig. S4. ChAdOx1 priming induces an innate and interferon response and amplifies the cytokine response to the subsequent MVA boost. (A)** Volcano plot showing differentially expressed genes following ChAdOx1 prime vaccination (24h) versus baseline in human peripheral blood. Differential expression from bulk RNA seq was assessed using edgeR *(65)*. Log2 fold changes (ChAdOx1 / baseline) and false discovery rates (FDR; Benjamini-Hochberg) were used for visualization. Genes with |log2FC| > 0.5 and FDR < 0.05 were considered differentially expressed. Selected innate immune and chemokine genes (CXCL10, IRF7, IFIH1, CCL2, CXCL9, CGAS, IL1B, CXCL11) are highlighted. **(B)** Mice were immunized with ChAd-P1A on day -7, one week later, they were boosted with MVA-P1A, and 6 hours after the serum was collected. **(C)** Comparison of cytokines fold-change 6 hours after ChAd+MVA or MVA alone. **(D)** Concentration of IFN$\gamma$ and (**E**) CXCL10 after each vaccination. **(F)** Gene Set Enrichment Analysis (GSEA) dot plot on the pseudobulk DESeq2 results described for Fig. 3C. Genes were ranked by their DESeq2 Wald statistic. GSEA was run using the fgsea package (v1.36.2) with 10,000 permutations. Gene sets from the MSigDB Hallmark collection (50 gene sets) and Reactome pathway database (C2:CP:REACTOME) were used. Gene sets with fewer than 15 or more than 500 members were excluded. Pathways with a positive normalized enrichment score (NES > 0, indicating upregulation at Day 1) and Benjamini-Hochberg adjusted p-value < 0.05 were retained. The top 10 pathways by NES are displayed separately for the Hallmark and Reactome collections as a bubble plot, with bubble size proportional to gene set size and color proportional to –log10(padj).

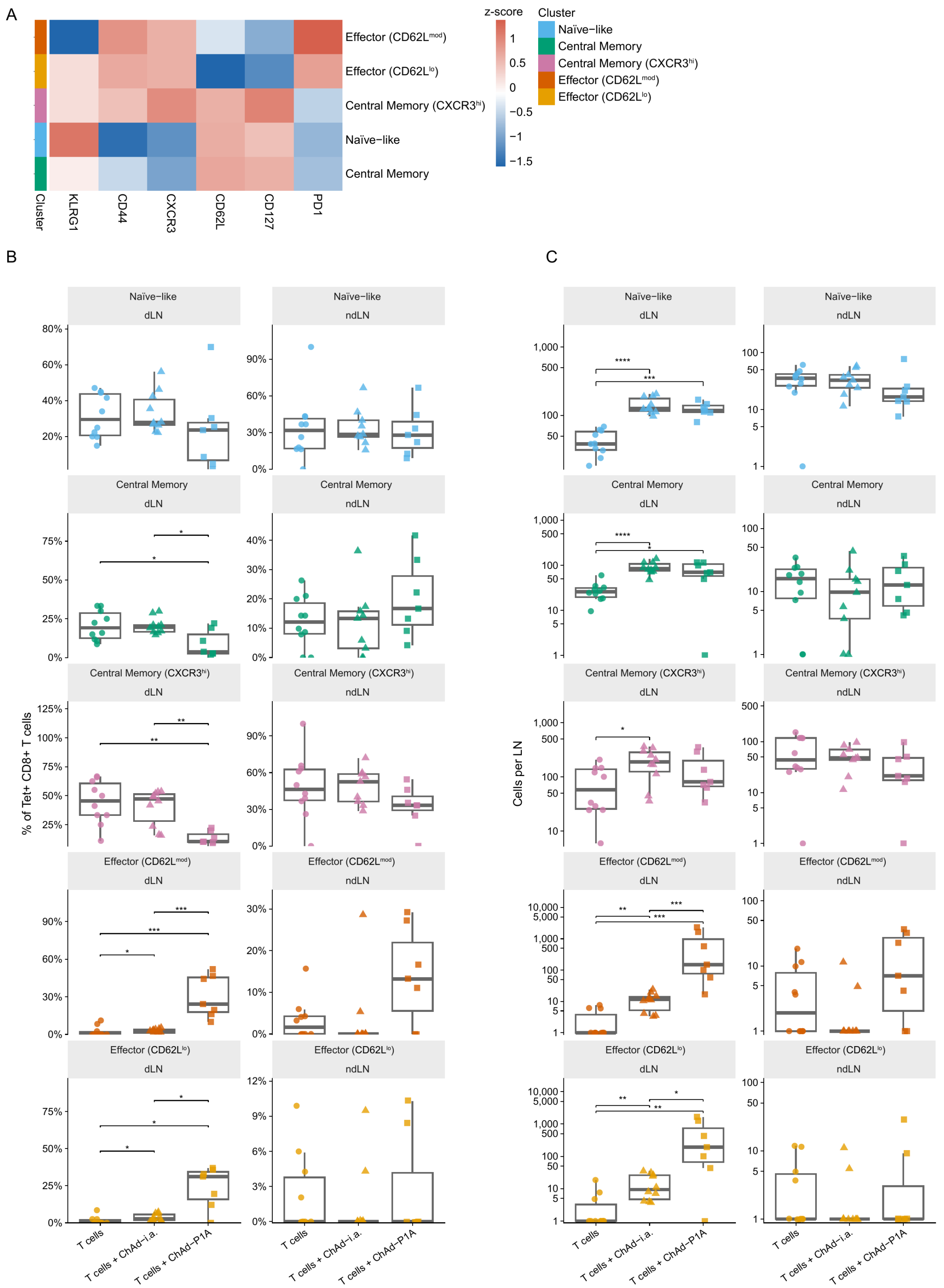

A
z-score
Cluster
Naïve−like
Central Memory
Central Memory (CXCR3hi)
Effector (CD62Lmod)
Effector (CD62Llo)
KLRG1
CD44
CXCR3
CD62L
CD127
PD1
B
C
dLN
ndLN
% of Tet+ CD8+ T cells
Cells per LN
T cells
T cells + ChAd−i.a.
T cells + ChAd−P1A

**Fig. S5**. **Phenotypic clustering of P1A-specific CD8$^+$ T cells in the draining and non-draining lymph nodes after ChAd vaccination.** (**A**) Heatmap (related to Fig. 3H) showing median arcsinh-transformed expression of each marker per cluster. Values are column-scaled (z-score) to highlight relative differences across clusters. Clusters (columns) were defined by k-means clustering (k = 5) applied to arcsinh-transformed flow cytometry data from Tet+ CD8+ T cells. Color scale: blue, below mean; white, mean; red, above mean. (**B**) Percentage and (**C**) total number of cells belonging to each of the clusters in the vaccine draining (dLN) and non-draining lymph nodes (ndLN). Statistical significance was assessed by Kruskal-Wallis test followed by pairwise Wilcoxon rank-sum tests with Benjamini-Hochberg correction; brackets indicate significant pairwise differences (p.adj $\leq$ 0.05).

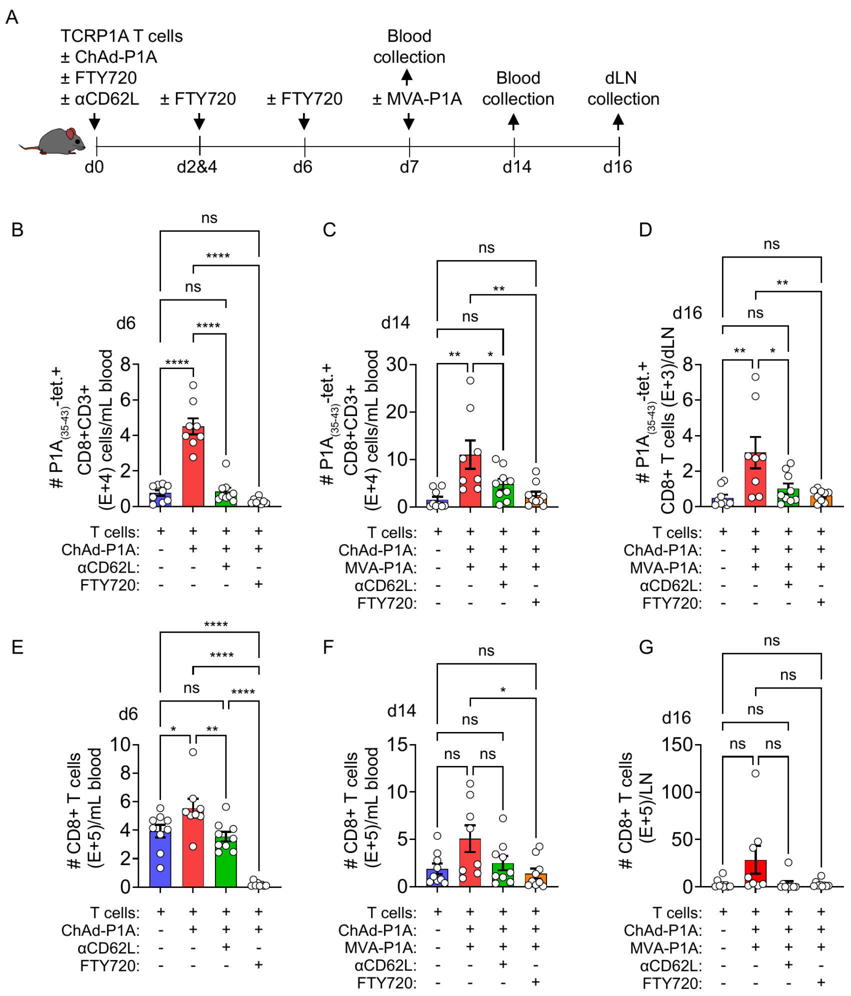

A
TCRP1A T cells
± ChAd-P1A
± FTY720
± αCD62L
± FTY720
± FTY720
Blood collection
± MVA-P1A
Blood collection
dLN collection
d0
d2&4
d6
d7
d14
d16
B
d6
### P1A(35-43)-tet.+ CD8+CD3+ (E+4) cells/mL blood
C
d14
### P1A(35-43)-tet.+ CD8+CD3+ (E+4) cells/mL blood
D
d16
### P1A(35-43)-tet.+ CD8+ T cells (E+3)/dLN
E
d6
### CD8+ T cells (E+5)/mL blood
F
d14
### CD8+ T cells (E+5)/mL blood
G
d16
### CD8+ T cells (E+5)/LN
T cells:
ChAd-P1A:
MVA-P1A:
αCD62L:
FTY720:

**Fig. S6**. **Disrupting T-cell homing to or egress from lymph nodes abrogates vaccine-mediated expansion of antigen-specific T cells.** (**A**) Mice were given both P1A T cells and ChAd-P1A on day 0. On day 7, mice received MVA-P1A. In addition to T cells and vaccines, some mice received *α*CD62L i.v. 2 hours before T cell injection to prevent T cell homing to LN or FTY720 i.p. 2-4 hours after T cell therapy (+ additional doses on days 2, 4 and 6) to prevent T cell exit from lymphoid tissues. Blood was collected on days 6 and 14, and dLNs on day 16 for flow cytometry analysis. **(B-D)** Concentration of P1A$_{(35\text{-}43)}$-tetramer+ CD8+ T cells in the blood on day 6 **(B)**, or day 14 **(C)**, or in the dLN on day 16 **(D)**. Concentration of CD8+ T cells in the blood on day 6 **(E)**, or day 14 **(F)**, or in the dLN on day 16 **(G)**.

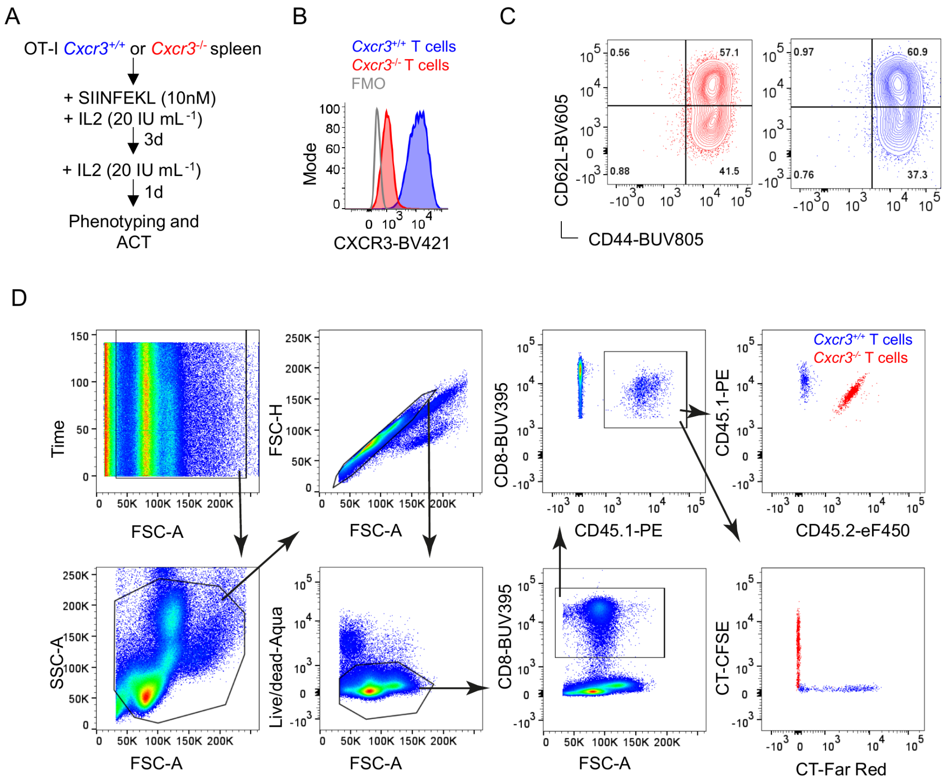
A
OT-I Cxcr3+/+ or Cxcr3-/- spleen
+ SIINFEKL (10nM)
+ IL2 (20 IU mL-1)
3d
+ IL2 (20 IU mL-1)
1d
Phenotyping and ACT
B
Cxcr3+/+ T cells
Cxcr3-/- T cells
FMO
Mode
CXCR3-BV421
C
CD62L-BV605
CD44-BUV805
0.56
57.1
0.88
41.5
0.97
60.9
0.76
37.3
D
Time
FSC-A
FSC-H
CD8-BUV395
CD45.1-PE
CD45.2-eF450
SSC-A
Live/dead-Aqua
CT-CFSE
CT-Far Red

**Fig. S7**. **Validation and phenotypic characterization of activated CXCR3-deficient and CXCR3-competent OT-I T cells. (A)** OT-I activation scheme. (**B**) Histogram with the CXCR3 expression after the activation protocol of CXCR3-deficient and competent OT-I cells. **(C)** Representative contour plots displaying the resulting CD62L and CD44 expression pattern. (**D**) Gating strategy for Fig. 4, B to F.

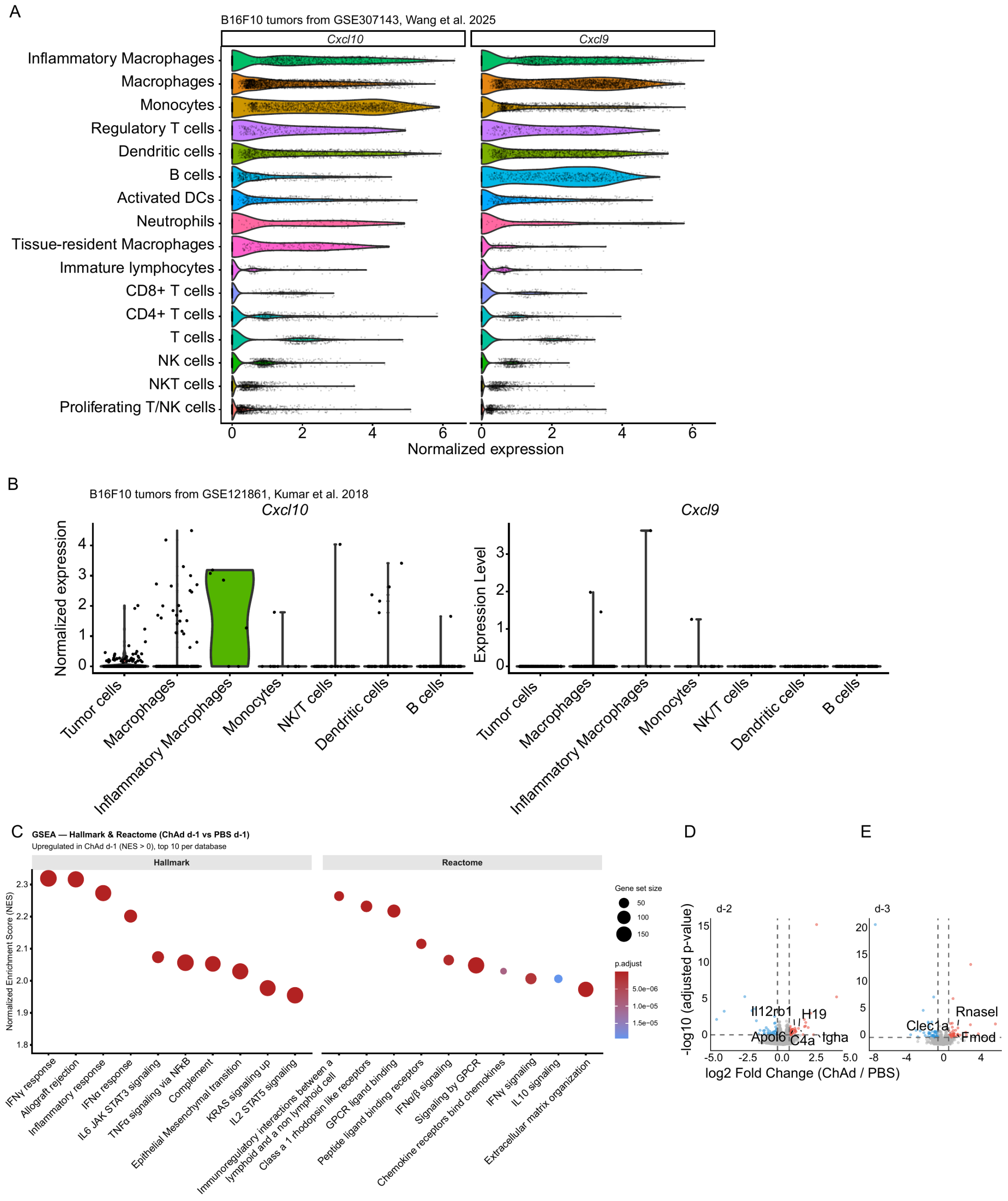
A
B16F10 tumors from GSE307143, Wang et al. 2025
Cxcl10
Cxcl9
Inflammatory Macrophages
Macrophages
Monocytes
Regulatory T cells
Dendritic cells
B cells
Activated DCs
Neutrophils
Tissue-resident Macrophages
Immature lymphocytes
CD8+ T cells
CD4+ T cells
T cells
NK cells
NKT cells
Proliferating T/NK cells
Normalized expression
B
B16F10 tumors from GSE121861, Kumar et al. 2018
Cxcl10
Cxcl9
Normalized expression
Expression Level
Tumor cells
Macrophages
Inflammatory Macrophages
Monocytes
NK/T cells
Dendritic cells
B cells
C
GSEA — Hallmark & Reactome (ChAd d-1 vs PBS d-1)
Upregulated in ChAd d-1 (NES > 0), top 10 per database
Hallmark
Reactome
Normalized Enrichment Score (NES)
Gene set size
p.adjust
IFNγ response
Allograft rejection
Inflammatory response
IFNα response
IL6 JAK STAT3 signaling
TNFα signaling via NFκB
Complement
Epithelial Mesenchymal transition
KRAS signaling up
IL2 STAT5 signaling
Immunoregulatory interactions between a lymphoid and a non lymphoid cell
Class a 1 rhodopsin like receptors
GPCR ligand binding
Peptide ligand binding receptors
IFNα/β signaling
Signaling by GPCR
Chemokine receptors bind chemokines
IFNγ signaling
IL10 signaling
Extracellular matrix organization
D
E
d-2
d-3
-log10 (adjusted p-value)
log2 Fold Change (ChAd / PBS)
Il12rb1
H19
Apol6
C4a
Igha
Rnasel
Clec1a
Fmod

**Fig. S8. Cellular sources of *Cxcl10* and *Cxcl9* in the B16F10 tumor microenvironment and the tumor transcriptional response to ChAdOx1. (A)** scRNA-seq data from B16F10 tumors (GSE307143) *(44)* were processed using Seurat (v5.1.0–5.4.0). After quality control (200–7,000 genes per cell; mitochondrial content < 25%), data were log-normalized, clustered using SNN-Louvain (20 PCs, resolution 0.4) and visualized by UMAP. Cell types were annotated manually based on canonical marker gene expression. *Cxcl10* and *Cxcl9* normalized expressions per annotated cell type are shown as violin plots. (**B**) A similar approach was done for GSE121861 *(45)*, which also consists of B16F10 tumor infiltrates. Cell types with fewer than 5 cells were excluded. (**C**) GSEA was performed on bulk RNA-seq data from B16F10-OVA tumors (ChAd vs PBS, day -1) using clusterProfiler *(82)*, with genes ranked by DESeq2 log2 fold change. MSigDB Hallmark and Reactome (C2:CP:REACTOME) gene sets curated for *Mus musculus* were tested (15–500 members; padj < 0.05, Benjamini-Hochberg). The top 10 upregulated pathways (NES > 0) per collection are shown as bubble plots (bubble size, gene set size; color, adjusted p-value). (D-E) Bulk RNA-seq was performed on B16F10-OVA tumors from mice treated with ChAdOx1 or PBS on days -2 (D) and -3 (E). Figure related to experimental design shown in Fig. 4G. Differential expression was assessed using DESeq2 (three biological replicates per group per timepoint); genes with |log2FC| > log2(1.5) and padj < 0.05 (Benjamini-Hochberg) were considered differentially expressed.

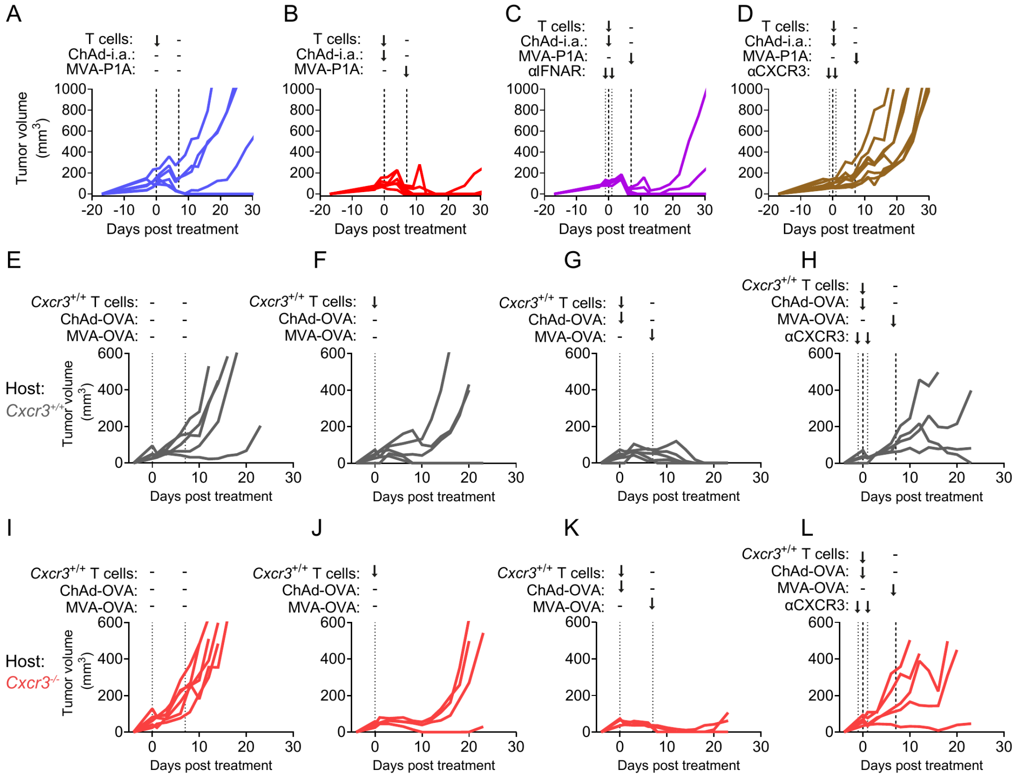
A
T cells:
ChAd-i.a.:
MVA-P1A:
Tumor volume (mm3)
Days post treatment
B
C
αIFNAR:
D
αCXCR3:
E
Cxcr3+/+ T cells:
ChAd-OVA:
MVA-OVA:
Host:
Cxcr3+/+
F
G
H
I
Cxcr3-/-
J
K
L

**Fig. S9. Individual tumor growth curves of mice treated with αCXCR3 or αIFNAR and of CXCR3-competent versus -deficient hosts.** (**A-D**) Individual tumor growth curves of mice shown in Fig. 5, A to C. (**E-L**) Individual tumor growth curves of experiment described in Fig. 5, D to I.

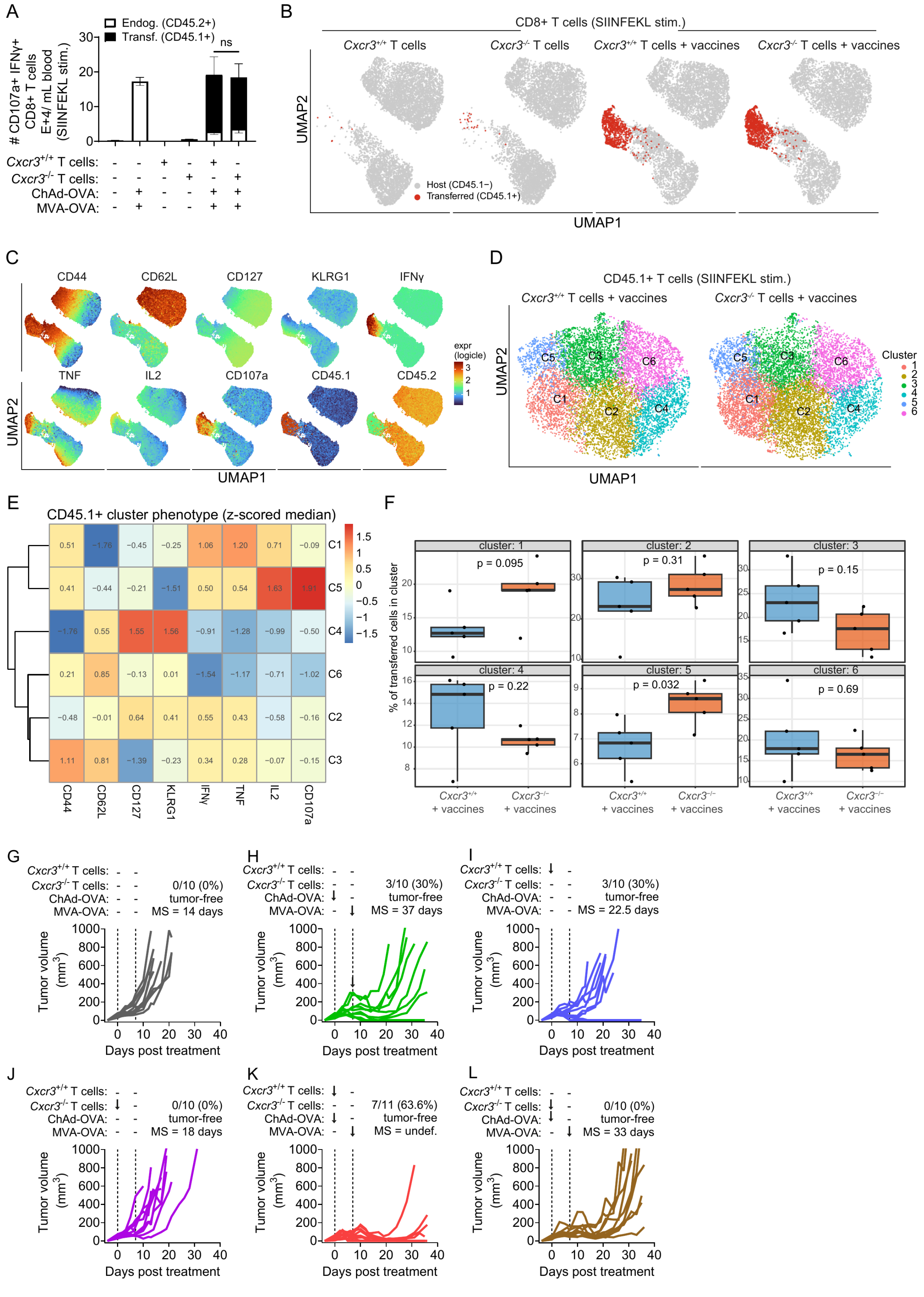
A
B
CD8+ T cells (SIINFEKL stim.)
Cxcr3+/+ T cells
Cxcr3-/- T cells
Cxcr3+/+ T cells + vaccines
Cxcr3-/- T cells + vaccines
Host (CD45.1−)
Transferred (CD45.1+)
UMAP1
UMAP2
C
D
CD45.1+ T cells (SIINFEKL stim.)
E
CD45.1+ cluster phenotype (z-scored median)
F
% of transferred cells in cluster
G
H
I
J
K
L
Days post treatment
Tumor volume (mm3)

**Fig. S10. CXCR3 is dispensable for the phenotype and effector differentiation of vaccine-primed antigen-specific CD8$^+$ T cells in blood.** (A) Bar graph displays the total number of circulating degranulating, IFNγ-producing (CD107a$^+$IFNγ$^+$) CD8$^+$ T cells after SIINFEKL peptide stimulation on day 13 after treatment of mice shown in Fig. 5J. Dark and white bars represent antigen-specific transferred (CD45.1$^+$) or endogenous (CD45.2$^+$) T cells, respectively. ns, two-sided Mann-Whitney U test. (B) UMAP of live CD3$^+$CD8$^+$ T cells from blood after SIINFEKL stimulation on day 13, concatenated per group and computed from CD44, CD62L, CD127, KLRG1, IFNγ, TNF, IL2 and CD107a (7,000 cells per group). Cells colored as host (CD45.1$^-$, gray) or transferred antigen-specific (CD45.1$^+$, red). Groups as in Fig. 5J. (C) Expression of the indicated markers overlaid on the UMAP in (B) (logicle scale; blue to red, low to high). (D) UMAP of transferred (CD45.1$^+$) T cells from the *Cxcr3*$^{+/+}$ and *Cxcr3*$^{-/-}$ + vaccines groups, clustered by k-means into six subpopulations (C1–C6) on the same eight markers. (E) Heatmap of per-cluster median expression (column z-scored) of each marker across the six clusters in (D); numbers indicate median (logicle) values. (F) Frequency of each cluster among transferred (CD45.1$^+$) T cells, *Cxcr3*$^{+/+}$ + vaccines vs *Cxcr3*$^{-/-}$ + vaccines (n = 5–6 mice per group). Each dot is one mouse; boxes show median and IQR. p, two-sided Mann-Whitney U test with Benjamini-Hochberg correction across clusters. (G–L) Individual tumor growth curves with proportion of tumor-free mice and median survival (MS) after treatment shown in Fig. 5J.

Table S1. Flow cytometry antibodies.

| Marker | Fluorochrome | Manufacturer | Staining step | Clone | Cat No: |
|---|---|---|---|---|---|
| CD3 | AF700 | Biolegend | Surface | 17A2 | 100216 |
| CD3 | BV421 | Biolegend | Intracellular | 17A2 | 100228 |
| CD8 | BUV395 | Invitrogen | Surface | 53-6.7 | 363-0081-82 |
| CD8 | BV421 | Biolegend | Surface | 53-6.7 | 100753 |
| CD8 | FITC | Biolegend | Surface | 53-6.7 | 100706 |
| CD8 | Spark UV 387 | Biolegend | Surface | 53-6.7 | 100798 |
| CD44 | APC-Cy7 | Biolegend | Surface | IM7 | 103028 |
| CD44 | BUV805 | BD Biosciences | Surface | IM7 | 741921 |
| CD44 | BV650 | Biolegend | Surface | IM7 | 103049 |
| CD45.1 | FITC | BD Biosciences | Surface | A20 | 553775 |
| CD45.1 | PE | Biolegend | Surface | A20 | 110707 |
| CD45.2 | eF450 | Invitrogen | Surface | 104 | 48-0454-82 |
| CD62L | APC-Cy7 | Biolegend | Surface | MEL-14 | 104427 |
| CD62L | BV605 | Biolegend | Surface | MEL-14 | 104412 |

| | | | | | |
|---|---|---|---|---|---|
| CD62L | BV650 | Biolegend | Surface | MEL-14 | 104453 |
| CD107a (LAMP1) | FITC | Biolegend | Surface together with peptide stimulation mix | 1D4B | 328605 |
| CD127 | PE-Cy5 | Biolegend | Surface | A7R34 | 135040 |
| CXCR3 | APC-Cy7 | Biolegend | Surface | CXCR3-173 | 126552 |
| CXCR3 | BV421 | Biolegend | Surface | CXCR3-173 | 126521 |
| CXCR3 | BV421 | Biolegend | Surface | S18001A | 155907 |
| Fixable Viability Dye (LD) | eFluor 506 | Thermo Scientific | Surface | - | 65-0866-18 |
| IFNγ | APC | Biolegend | Intracellular | XMG1.2 | 505810 |
| IL2 | APC-Cy7 | BD Biosciences | Intracellular | JES6-5H4 | 560547 |
| IL2 | PE | Biolegend | Intracellular | JES6-5H4 | 503808 |
| KLRG1 | APC-eF780 | Biolegend | Surface | 2F1/ KLRG1 | 138425 |
| KLRG1 | BV421 | BD Biosciences | Surface | 2F1 | 562897 |
| PD1 | PE-Cy7 | Biolegend | Surface | 29F.1A12 | 135216 |
| TNFα | BV650 | Biolegend | Intracellular | MP6-XT22 | 506333 |

Table S2. Surface antibody panel used for flow cytometric cell sorting before single-cell RNA-sequencing.

| Marker | Fluorochrome | Manufacturer | Staining step | Clone | Cat No: |
|---|---|---|---|---|---|
| CD45 | PE | BioLegend | Surface | 30-F11 | 103105 |
| CD3ε | BV785 | BioLegend | Surface | 145-2C11 | 100355 |
| CD4 | BV650 | BioLegend | Surface | RM4-5 | 100545 |
| CD8α | APC | BioLegend | Surface | 53-6.7 | 100711 |
| CD44 | BV421 | BioLegend | Surface | IM7 | 103039 |
| CD62L | PE-Cy7 | BioLegend | Surface | MEL-14 | 104417 |
| B220 | APC-Cy7 | BioLegend | Surface | RA3-6B2 | 103223 |
| IgM | BV605 | BioLegend | Surface | RMM-1 | 406523 |
| IgD | BV711 | BioLegend | Surface | 11-26c.2a | 405731 |
